# A Validated Functional Analysis of PALB2 Missense Variants for Use in Clinical Variant Interpretation

**DOI:** 10.1101/2020.08.27.270553

**Authors:** Sarah E. Brnich, Eyla Cristina Arteaga, Yueting Wang, Xianming Tan, Jonathan S. Berg

## Abstract

Clinical genetic testing readily detects germline genetic variants. Yet, the evidence available for variant classification as benign or pathogenic is often limited by the rarity of individual variants, leading to many “variant of uncertain significance” (VUS) classifications. VUS cannot guide clinical decisions, complicating counseling and management. Laboratory assays can potentially aid reclassification, but require benchmarking against variants with definitive interpretations to have sufficient predictive power for clinical use. Of all clinically identified germline variants in hereditary breast cancer gene *PALB2* (Partner and Localizer of BRCA2), ~50% are VUS and ~90% of VUS are missense. Truncating PALB2 variants have homologous recombination (HR) defects and instead rely on error-prone non-homologous end-joining (NHEJ) for DNA damage repair (DDR). Recent reports show some missense PALB2 variants may also be damaging, but thus far functional studies have lacked benchmarking controls. Using the Traffic Light Reporter (TLR) to quantify cellular HR and NHEJ using fluorescent markers, we assessed variant-level DDR capacity in hereditary breast cancer genes. We first determined the TLR’s dynamic range using *BRCA2* missense variants of known significance as benchmarks for normal/abnormal HR function. We then tested 37 *PALB2* variants, generating functional data for germline *PALB2* variants at a moderate level of evidence for a pathogenic interpretation (PS3_moderate) for 8 variants, or a supporting level of evidence in favor of a benign interpretation (BS3_supporting) for 13 variants, based on the ability of the assay to correctly classify *PALB2* validation controls. This new data can be applied in subsequent variant interpretations for direct clinical benefit.

## INTRODUCTION

Detection of germline genetic variants has increased with the rise of clinical genetic testing.^1–3^ Yet, the rarity of many variants limits the evidence available to determine if a variant is pathogenic or benign, leading to many “variant of uncertain significance” (VUS) classifications. VUS cannot guide clinical decisions, complicating post-test patient counseling and management.^4–6^ Laboratory tests of a variant’s functional impact can potentially aid reclassification^4,7^, but require benchmarking against variants with definitive interpretations to have sufficient predictive power for clinical use.^8^

Even among genes with clear links to disease and well-understood functions, VUS account for a large amount of total identified variation. For example, many of the genes implicated in hereditary breast cancer (MIM: 114480) have known roles in homology-directed repair (HR) of DNA double-strand breaks (DSB) under normal conditions, but the clinical interpretation of missense variation in these genes remains a challenge. In one such gene, *PALB2* (Partner and Localizer of BRCA2 [MIM: 610355]), ~50% of all clinically identified variants in the Clinical Variant database (ClinVar) are VUS and ~90% of VUS are missense variants.^9^

For high-fidelity repair of DSBs via HR, the coiled-coil (CC) regions in BRCA1 and PALB2 must first interact to stably localize PALB2 to the sites of DNA double-strand breaks.^10,11^ In turn, the PALB2 WD40 domain interacts with BRCA2 and RAD51 to form a complex directing the use of a homologous template strand to correctly repair DSBs.^11–14^ As such, PALB2 acts as the “molecular scaffold” in the BRCA1-PALB2-BRCA2 complex.^10^ PALB2 homodimers can also form through self-interaction of the CC domain, preventing heterodimerization with BRCA1 and potentially regulating HR efficiency.^15,16^ Defects in HR can lead to repair via alternative non-homologous end-joining (alt-NHEJ), an error-prone mechanism that leads to small insertions and deletions.^17–19^

Truncating (nonsense and frameshift) variants comprise the majority of pathogenic variation in *PALB2* and demonstrate HR defects, relying instead on error-prone alt-NHEJ for DNA damage repair (DDR). As such, tumors with germline *PALB2, BRCA1* (MIM: 113705), and *BRCA2* (MIM: 600185) defects demonstrate a pattern of genome-wide instability and a higher mutational burden.^20,21^ Recent reports have indicated that some missense PALB2 variants may also be damaging, but thus far functional studies have lacked sufficient benchmarking controls for validated use in clinical variant interpretation.^22–25^ Other hereditary cancer genes in the HR pathway have been sufficiently well-studied and have good benchmarking controls. For example, BRCA2 missense variants have been characterized previously by their capacity for HR using the direct repeat-GFP (DR-GFP) reporter assay, where GFP serves as a marker for HR following an induced DNA DSB.^26,27^ Benign variants retain HR capacity and thus strongly express GFP, while pathogenic variants defective for HR do not. The established BRCA2 benign and pathogenic variants used in developing this assay constitute a gold standard panel for functional assay validation.^27,28^ We evaluated whether we could use these same BRCA2 gold standard missense variants to establish an assay of DDR for PALB2 missense variants.

The American College of Medical Genetics and Genomics (ACMG)/Association for Molecular Pathology (AMP) sequence variant interpretation guidelines outline each evidence criterion and the assigned weight (supporting, moderate, strong, very strong) at which the evidence can be applied.^1^ These categorical strengths have also been modeled as Bayesian Odds of Pathogenicity (OddsPath).^29^ Together with the ClinGen Sequence Variant Interpretation Working Group (SVI), we recently described recommendations for using statistical analyses to determine the strength at which the ACMG/AMP functional evidence criteria can be applied in support of a pathogenic or benign classification (PS3 or BS3 codes, respectively).^8^ Adhering to this, we sought to derive the strength of evidence that could be applied for PALB2 clinical variant interpretation, based on the performance of the assay in PALB2 variants with known clinical relevance.

## MATERIAL AND METHODS

### Plasmids, cloning, and siRNA

Human N-terminal, FLAG-tagged *PALB2* was a gift from Daniel Durocher (pDEST-FRT/T0-FLAG-PALB2, plasmid #71114, Addgene)^30^. Human wild-type (WT) *BRCA2* was obtained from pcDNA3 236HSC WT (BRCA2), a gift from Mien-Chie Hung (#16246, Addgene)^31^. *BRCA2* cDNA was then Gateway cloned into pDONR221 (kind gift of Jeff Sekelsky). The *attB* sites were introduced by touchdown, gradient polymerase chain reaction (PCR), followed by stitching PCR and restriction digest/ligation using *Acl-I* and *SphI-HF* sites to generate a complete *att*B-PCR product. BP recombination reaction was performed according to manufacturer protocol (Thermo Fisher Scientific). To standardize plasmid delivery across genes, pDONR221-BRCA2 was then subcloned into pDEST-FRT/T0-FLAG by LR reaction, according to manufacturer protocol (Thermo Fisher Scientific).

The *BRCA2* and *PALB2* constructs were made siRNA resistant by introducing three silent mutations with the QuikChange II XL Site-Directed Mutagenesis Kit (Agilent), according to the manufacturer’s protocol. For *BRCA2*, siRNA resistance was conferred with the following silent mutations: c.4773T>C, c.4779A>G, c.4785G>A. For *PALB2*, siRNA resistance was achieved with the following silent mutations: c.1635A>G, c.1641C>A, c.1647C>T. All variants were generated in the siRNA-resistant constructs using the same site-directed mutagenesis kit as above.

Constructs were confirmed by Sanger sequencing (Genewiz). siRNA target sequences and sequences of primers used for cloning, site-directed mutagenesis, and Sanger confirmation are provided as supplemental material (Tables S1-S6).

### Cell culture and Traffic Light Reporter (TLR) cell line generation

HEK293T/17 cells were purchased from American Type Culture Collection (ATCC) and tested negative for mycoplasma. For TLR lentivirus production, we used Lipofectamine 2000 (Invitrogen) to transfect HEK293T/17 cells with the pCVL Traffic Light Reporter (TLR) construct, a gift from Andrew Scharenberg (#31482, Addgene)^32^, and lentiviral packaging plasmids pMD2.G and pCMV delta R8.2, gifts of Didier Trono (Addgene plasmids #12259 and 12263).^33^ HEK293T/17 cells were transduced at a low multiplicity of infection (100 colony-forming units) with TLR lentivirus to generate a stable, polyclonal cell line with a high likelihood of single integration, as previously described.^33^ We will henceforth refer to these cells as 293T/TLR.

293T/TLR cells were maintained in Dulbecco’s Modified Eagle’s Medium-High Glucose (DMEM-H) (Sigma-Aldrich) with 10% fetal bovine serum (FBS) (VWR,), 1% antibiotic-antimycotic (Gibco), 4mM L-Glutamine (Corning), and 10 μg/mL puromycin (Corning). Cells were grown at 37°C, 5% CO_2_ in a humidified incubator. All siRNA and DNA transfections were performed in antibiotic-free DMEM-H with 10% FBS and 4 mM L-Glutamine.

### Preliminary assay validation

After the generation of our stable 293T/TLR cell line, we performed a preliminary test of the system, as previously described.^32^ On day one, we seeded 6 x 10^5^ 293T/TLR cells per well of a 12-well plate and transiently transfected them with 1.0 μg of a homologous donor template pGFPdonor-BFP (pRRL SFFV d20GFP.T2A.mTagBFP Donor, Addgene plasmid #31485)^32^, I-Sce1 nuclease pI-SceI-IFP (pRRL sEF1a HA.NLS.Sce(opt).T2A.IFP, Addgene plasmid #31484)^32^, or both using Lipofectamine 2000 (Invitrogen). Antibiotic-free media was replaced the next day. On day three, cells were transferred to a 6-well plate. On day four, 1 x 10^6^ cells were analyzed by flow cytometry using an LSRFortessa (BD Biosciences).

### Traffic Light Reporter Assay analysis of *BRCA2* and *PALB2* variants

For analysis of DDR outcomes with BRCA2 or PALB2 variants, experiments were performed as follows. On day one, we reverse transfected approximately 3.0 x 10^5^ - 5.0 x 10^5^ 293T/TLR cells with 75 nM of siRNA per well of a 6-well plate using Lipofectamine RNAiMAX (Invitrogen) according to the manufacturer’s protocol. Twenty-four hours later, medium was changed and Lipofectamine 2000 was used to forward transfect cells with 1.25 μg of the IFP-tagged I-SceI nuclease construct, 1.25 μg of pGFPdonor-BFP, and either 1.25 μg of empty vector (EV), 1.25 μg of the indicated siRNA-resistant pDEST-FLAG/BRCA2, or 0.71 μg of the indicated siRNA-resistant pDEST-FLAG/PALB2 construct. 72 hours after DNA transfection, cells were harvested by enzymatic release, resuspended in cold 1% Bovine Serum Albumin (BSA) (Sigma-Aldrich) in Dulbecco’s Phosphate-Buffered Saline (D-PBS) (Gibco) to 1 x 10^6^ cells/mL, and 1.0-3.0 x 10^6^ cells were filtered for analysis by flow cytometry. Remaining cells were pelleted and stored at −20°C for subsequent RNA analysis.

Flow cytometry analysis was performed using a LSRFortessa cell analyzer (BD Biosciences). Live cells were gated based on forward scatter area versus side scatter area. Single cells were then gated based on forward scatter area versus forward scatter height. GFP was excited by a 488-nm laser and detected by a 530/30 filter. mCherry was excited by a 561-nm laser and detected by a 610/20 filter. Infrared Protein (IFP) was excited by a 640-nm laser and detected by a 710/50 filter. Blue Fluorescent Protein (BFP) was excited by a 405-nm laser and detected by a 450/50 filter. Data were analyzed using FACS Diva (BD Biosciences) and FCS Express 6 (De Novo Software) software.

At least 10,000 doubly transfected (IFP^+^/BFP^+^), single, live cells were analyzed per condition. From this population of cells, we obtained readouts of GFP^+^/mCherry^+^ ratio (HR/NHEJ) and the percent of cells that were GFP^+^ (an indicator of %HR). Each variant was tested in at least three independent experiments. Every 6-well plate included the following conditions as controls: non-targeting siRNA (siNT)-treated cells rescued with EV, siBRCA2- or siPALB2-treated cells rescued with EV, and siBRCA2- or siPALB2-treated cells rescued with siRNA-resistant WT *BRCA2* or *PALB2*.

### Western blots

Cell pellets were resuspended in Pierce RIPA buffer (Thermo Scientific), containing 1mM Phenylmethylsulfonyl fluoride (PMSF) (Thermo Scientific), 1X complete protease inhibitor cocktail (Roche Diagnostics), 0.1% 2-Mercaptoethanol (Gibco), 0.1% Dithiothreitol (DTT) (Thermo Scientific), and 1 tablet of Pierce phosphatase inhibitor (Thermo Scientific) per 10 mL of buffer. Resuspended pellets were shaken for 10 minutes at 4°C and lysed by sonication in a 4°C water bath (Diagenode Biorupter) for five minutes (30 seconds on/30 seconds off). Lysate was then centrifuged for 10 minutes at 14,000 x *g* at 4°C and supernatant transferred to a new tube. Protein concentration was measured by reducing-agent compatible microplate BCA Assay (Thermo Scientific) according to kit protocol. Plates were read using a GloMax Multi-Detection System (Promega).

For western blotting, reduced whole cell lysates were electrophoresed on a NuPAGE 7% Tris-Acetate pre-cast 1.0 mm gels (Invitrogen) for 50 minutes at 150 V and transferred to polyvinylidene difluoride (PVDF) membrane (Merck Millipore) at 30 V for 1 hour. Membranes were blocked in 5% BSA in Tris-buffered saline (TBS) (Santa Cruz Biotechnology, Inc.) for 1 hour at room temperature. Primary antibodies used were a mouse monoclonal anti-FLAG antibody (1:1,000; #F1804, Sigma-Aldrich), a polyclonal rabbit antibody against an epitope between residues 200-250 of PALB2 (1:2,000; #A301-247A, Bethyl Laboratories), and a rabbit monoclonal antibody against GAPDH (1:2,500; #AB9485, Abcam). Primary antibody incubation was performed for 1 hour at room temperature, or overnight at 4°C. Secondary antibodies used were IRDye 800CW Goat anti-mouse or anti-rabbit IgG (1:20,000-1:25,000, #925-32210 and 925-32211, respectively) (LI-COR). Membranes were incubated with secondary antibodies for 1 hour at room temperature, protected from light. After primary and secondary antibody incubation, membranes were washed 3 times for 10 minutes at room temperature with 0.1% Tween 20 (Sigma-Aldrich) in 1X TBS. Membranes were then rinsed with 1X TBS and blots were visualized using a LI-COR Odyssey CLx Imager.

### Real-Time PCR (RT-PCR)

RNA was isolated and purified from pelleted 293T/TLR cells using a RNeasy Plus Mini Kit (Qiagen). 100 ng of RNA was used as template with a TaqMan RNA-to-C_T_ 1-Step Kit (Applied Biosystems) for gene expression analysis of *PALB2* or *BRCA2* relative to the beta-2-microglobulin (*B2M*) gene (MIM: 109700) in 25 mL triplicate reactions run on an Applied Biosystems StepOne Real-Time PCR System. TaqMan Gene Expression Assay IDs (Applied Biosystems) used were Hs00609073_m1 (*BRCA2*), Hs00954121_m1 (*PALB2*), and Hs00187842_m1 (*B2M*).

### Normalization

For the BRCA2 and PALB2 TLR assays, flow cytometry data from each experimental condition was normalized to the in-plate mock control (siNT + EV). Then, we utilized the assay results for EV and WT controls, which are included on each plate, to estimate a plate effect using a linear mixed model (LMM), which assumes in-plate effects as fixed effects and between-plate effects as random effects. We used the lme function from the nlme package in R^34^ to estimate the plate effects and implement the normalization, i.e., subtracting corresponding estimated plate effects from all reads. Assay results for BRCA2 variants were adjusted based on LMM with random effects normalization using BRCA2 WT and EV controls, and assay results for PALB2 variants were separately adjusted based on LMM with random effects normalization using PALB2 WT and EV controls.

### Gaussian Mixture Model

We used the normalized HR/NHEJ readout for WT and EV controls to generate Gaussian mixture models (GMM) for BRCA2 and PALB2, respectively, using the `normalmixEM` function from the `mixtools`^35^ package in R^34^, setting k to 2. This function provides as output the mean, standard deviation and mixture weight for each estimated distribution, and provides the posterior probability for a given input value to belong to each distribution. Then, using the dnorm function we are able to calculate the probability of any arbitrary assay read-out value to belong to either of the distributions (based on the mean and standard deviation from the GMM). We can then calculate the ratio of these probabilities and locate the threshold at which there is a 10:1 ratio in favor of belonging to either the abnormal or normal distribution.

### Regression analysis

Scatterplots and linear regression were generated to test the correlation between functional data and in silico predictors, using squared correlation coefficients (R^2^) to measure the strength of correlation (GraphPad Prism v8.3.0). Three studies of the functional impact of PALB2 missense variants were recently published.^23–25^ To test the correlation between the TLR’s HR/NHEJ readout and recently published HR functional data, we repeated our regression analysis using published %HR efficiency values^25^ or homology-directed repair (HDR) fold change^23^. We cannot directly compare HR/NHEJ to HR readout from the third study^24^, as raw HR data was not published.

## RESULTS

### Establishment of an *in vivo* cellular assay to detect HR/NHEJ outcomes

Originally developed for optimization of genome editing protocols, the Traffic Light Reporter (TLR) uses cellular GFP and mCherry expression to visualize the repair of induced DNA DSBs by HR or alt-NHEJ, respectively (Figure 1a).^32^ We established this assay system in HEK293T/17 cells and demonstrated that it can serve as a reliable read-out for HR and NHEJ outcomes in our hands (Figure 1b). Untreated 293T/TLR cells are non-fluorescent at baseline, since the stably integrated GFP construct is disrupted by an I-SceI nuclease target site, and mCherry expression is kept out of reading frame by a T2A linker. Transfection with a plasmid containing a truncated GFPdonor (*truncGFP*) can be monitored by BFP expression, while I-SceI nuclease transfection can be monitored by IFP expression. When I-SceI is expressed, it can induce a DSB within the TLR GFP construct, and in the absence of a homologous repair template, alt-NHEJ repair events are detected by mCherry expression when a two-base pair frameshift occurs (about one-third of events)^32^. Co-transfection of both I-SceI and *truncGFP* donor template results in DSBs that are repaired by either HR or alt-NHEJ, indicated by GFP and mCherry expression, respectively.

**Figure 1.**
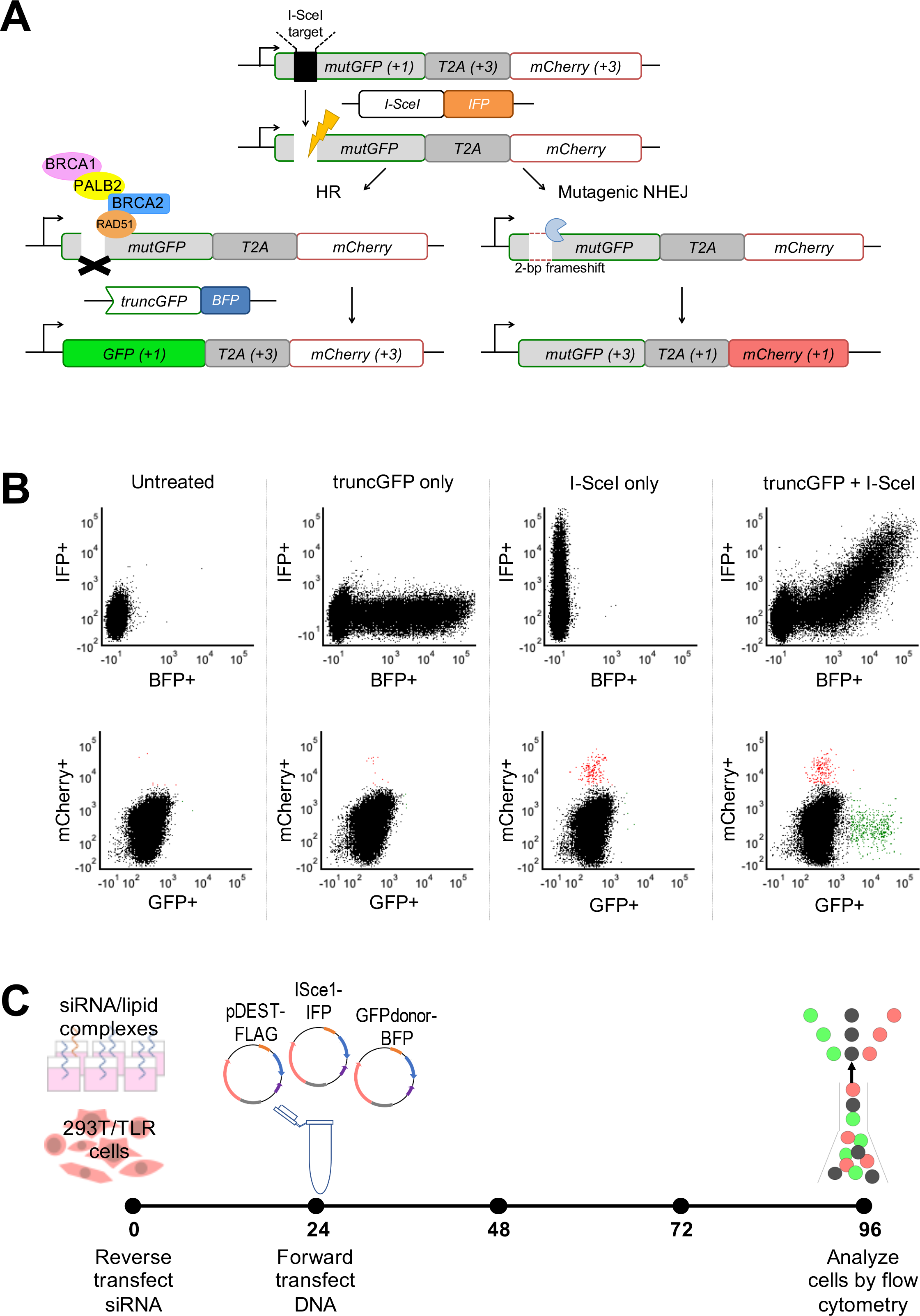
Traffic Light Reporter assay schematic, flow cytometry output, and final assay protocol. (A) The TLR assay, adapted from Certo et al.^32^, utilizes a cell line with a stably integrated construct encoding mutGFP (a mutant version of the GFP cDNA that does not produce green fluorescent protein at baseline due to an engineered I-SceI nuclease target site) connected via the T2A linker to an out-of-frame mCherry cDNA. Position relative to reading frame is indicated in parentheses, where +1 leads to gene expression, while +3 is 2 base pairs (bp) out of reading frame. A DNA double-strand break in mutGFP is induced by transfection with a plasmid encoding IFP and I-SceI nuclease. A plasmid encoding BFP and a truncated GFP construct (truncGFP) provides a homologous donor sequence that can be used by the cell to repair the DNA double-strand break via the homology-directed repair (HR) pathway (left side). However, if the donor sequence is absent, or if HR is not intact (right side), the cell can undergo mutagenic non-homologous end-joining (NHEJ) thereby restoring mCherry expression if it results in a 2-bp frameshift (about one-third of events). (B) Flow cytometry plots show mean fluorescence intensity of live, single cells. Expression of IFP (indicating presence of the I-SceI plasmid) versus BFP (indicating presence of the truncGFP donor plasmid) is shown in the upper panels. Expression of GFP (indicating successful HR) versus mCherry (indicating NHEJ) is shown on the lower panels. Untreated 293T/TLR cells are negative for BFP, IFP, GFP, or mCherry. Cells transfected only with truncGFP donor plasmid are reflected by a BFP positive population, otherwise negative for IFP, GFP, or mCherry. Cells transfected with I-SceI nuclease are IFP positive, and some of the population also expresses mCherry, indicating DSB repair by NHEJ due to absence of the homologous donor template. Cells transfected with all components of the TLR assay exhibit presence of both truncGFP and I-SceI (double positive BFP/IFP population), with distinct populations positive for mCherry or GFP indicating DSB repair by NHEJ or HR, respectively). (C) Optimized timing of the TLR assay used for siRNA knock-down and re-expression of BRCA2 and PALB2 variants in 293T/TLR cells. Stably transfected 293T/TLR cells are reverse transfected with siRNA at time 0, followed by transient co-transfection of rescue plasmid (pDEST-FLAG), I-SceI nuclease (co-expressing IFP) and truncGFP donor construct (co-expressing BFP) 24 hours later. Flow cytometry is performed 72 hours after DNA transfection to detect DDR outcomes.

We optimized TLR assay timing and siRNA knockdown efficiency (see Supplemental Materials and Methods for details) in order to determine a standardized assay protocol (Figure 1c) for use in all subsequent validation experiments. Briefly, we obtained ~70% siRNA-mediated knockdown of the target gene (*BRCA2* or *PALB2*) relative to siNT-treated cells at 24-hours (Figure S1b) and >50% knockdown at the time of flow cytometry (96-hours post-siRNA transfection) (Figure S2), with concomitant decrease in protein expression demonstrated by western blot (Figure S1c). Expression of I-SceI and *truncGFP* plasmids peaked between 24-72 hours post-transfection, and GFP or mCherry repair events can be detected after 48-72 hours (Figure S1a). Expression of FLAG-tagged BRCA2 or PALB2 protein from the rescue plasmid was detectable by western blot by 24 hours after transfection (Figure S1c). Therefore, using the experimental timing as shown in Figure 1c, we anticipate that reduction of endogenous BRCA2 or PALB2 protein expression, induction of DSB via I-SceI expression, and reconstitution via the rescue plasmid, will occur during a relevant time window for detecting the impact of missense variants on DDR outcomes as indicated by changes in the proportion of cells that are GFP or mCherry positive.

### Initial evaluation of the TLR assay using gold-standard *BRCA2* validation controls

In order to establish the TLR assay for use in clinical variant interpretation, we utilized a set of “gold standard” missense variants in *BRCA2* (8 pathogenic and 8 benign) that can be confidently interpreted without functional evidence and have previously been used as validation controls for a similar HR assay.^27,28^ Cells were transfected with either EV or siRNA-resistant FLAG-BRCA2 (encoding either WT, benign, or pathogenic BRCA2 variants) in a 6-well plate format. We measured the ratio of GFP^+^/mCherry^+^ cells (HR/NHEJ) for at least 10,000 doubly transfected (IFP^+^/BFP^+^) cells for each condition and examined each BRCA2 variant in at least three independent experiments.

In this 6-well plate implementation of the TLR assay, three wells of each plate were dedicated to controls: mock treatment consisting of siNT-treated cells rescued with EV, siBRCA2-treated cells rescued with WT BRCA2 cDNA “normal” control, and siBRCA2-treated cells rescued with EV “abnormal” control. Therefore, experimental replicates for each of the variants were distributed across numerous batches processed over time in different plates. This set-up could lead to variability between plates based on subtle factors such as the cell passage number, confluency, cell cycle phase; media and/or reagent batches; siRNA knockdown and/or transient re-expression; incubator temperature and/or CO_2_ concentration; variation in timing of experimental manipulations; or other unknown variables. To address this potential source of irregularity, we first adjusted the raw %HR and HR/NHEJ ratio within each plate by normalizing to the mock treated well. We then performed additional normalization across the plates using a linear mixed model (LMM) with random effects in order to adjust for factors that could change over time. While the raw data already demonstrated a relatively clear separation between the classes (Figure 2a), LMM random effect normalization improved this separation dramatically (Figure 2b).

**Figure 2.**
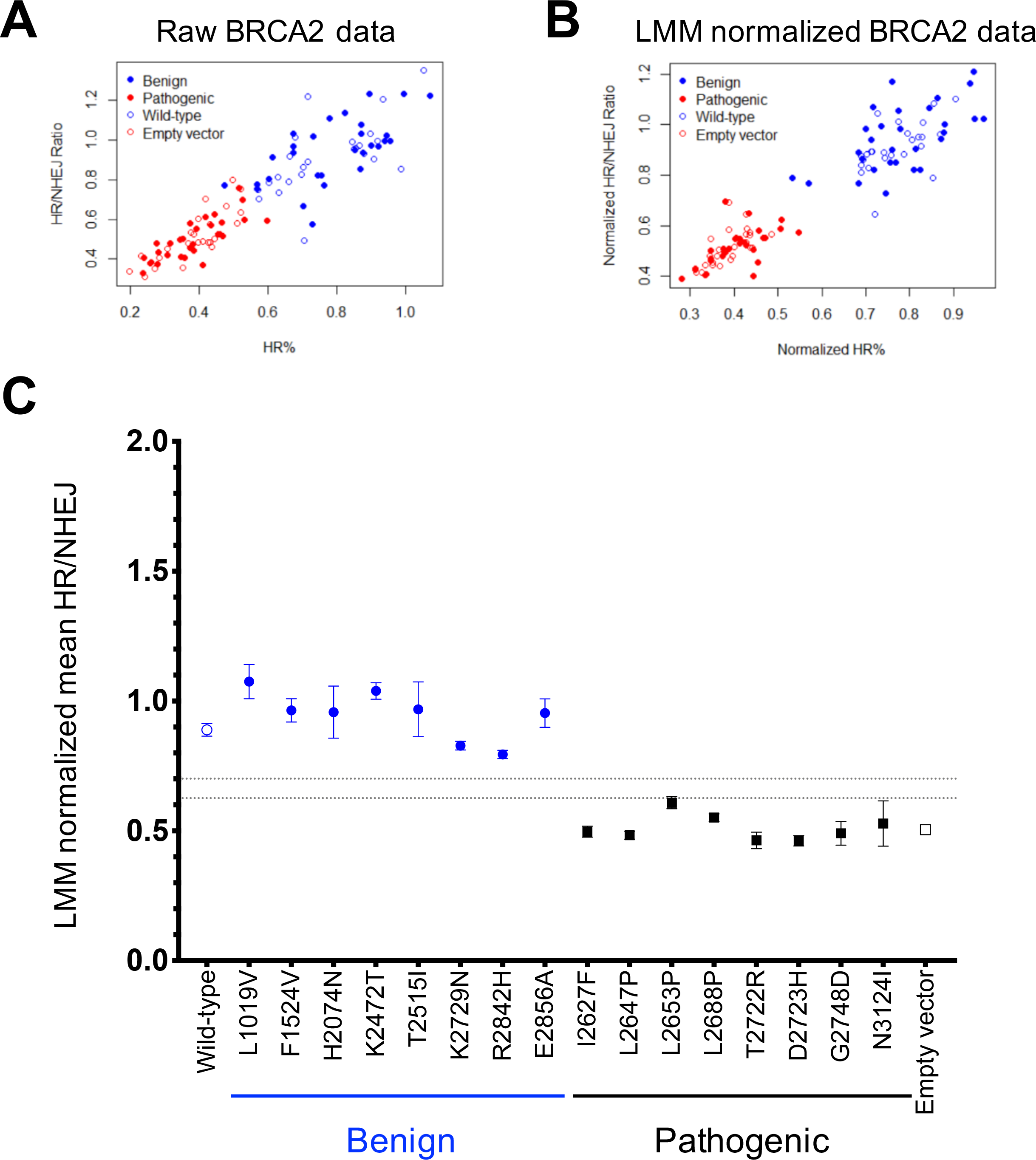
Validation of the TLR assay using BRCA2 gold-standard controls. (A) Raw flow cytometry data for %HR and HR/NHEJ ratio are shown for BRCA2 wild-type, empty vector, and gold-standard benign and pathogenic validation controls (normalized against the internal “mock” treated control for each plate). (B) Batch effects due to plate-to-plate experimental variability were addressed by normalization based on a Linear Mixed Model (LMM) with random effects, utilizing the assay results for wild-type and empty vector controls included in each plate, demonstrating improvement in separation between the different classes of variants. (C) TLR assay results for the BRCA2 experiments are summarized, demonstrating HR function after siBRCA2 knockdown and rescue with plasmids containing BRCA2 variants or controls. HR function is rescued by BRCA2 wild-type cDNA and benign validation controls but is impaired with empty vector and pathogenic validation controls. Each rescue condition is depicted across the x-axis. The y-axis represents the LMM normalized HR/NHEJ ratio as the mean ± SEM of at least three independent experiments. For wild-type and empty vector controls, N=21. Thresholds for normal and abnormal functional impact are indicated by gray dashed lines.

WT and benign *BRCA2* variants restore HR capacity as indicated by a higher HR/NHEJ ratio (means range from 0.794 – 1.075) while EV and pathogenic *BRCA2* variants demonstrate abnormal HR function, as indicated by a lower HR/NHEJ ratio (means range from 0.462 – 0.609). Although there is a clear visual separation between benign and pathogenic variant control groups (Figure 2c), the goal of this instance of the TLR assay is to provide functional evidence that can be used for clinical variant interpretation according to the ACMG/AMP guidelines^1^ with assignment of the strength of evidence according to a Bayesian formulation of the combining rules^29^ supported by statistical validation based on odds of pathogenicity^8^. In order to define thresholds that define “normal,” “abnormal,” and “indeterminate” TLR assay read-outs, we therefore utilized an unsupervised Gaussian mixture model (GMM) to estimate the density distribution contributions for the WT and EV controls. The distribution that corresponds to WT controls had a mean HR/NHEJ of 0.894 (standard deviation 0.098) and the distribution that corresponds to EV controls had a mean HR/NHEJ of 0.501 (standard deviation 0.066). We then defined thresholds based on a 10:1 ratio of the probability to belong to either the normal (HR/NHEJ > 0.701) or abnormal (HR/NHEJ < 0.627) distributions (Figure S3). Using these thresholds, all 8 benign variants and 7 out of 8 pathogenic variants would be correctly classified, with 1 pathogenic control (L2653P) classified as intermediate or indeterminate. Therefore, using the odds of pathogenicity (OddsPath) levels promulgated by Tavtigian et al.^29^ and further elaborated in Brnich et al.^8^, this functional evidence would reach moderate strength for PS3 and moderate strength for BS3 (albeit with caveats about “normal” function in this assay not necessarily reflecting all potential functional read-outs). With additional pathogenic and benign controls, we anticipate that the strength of the assay could be improved. Nevertheless, these results demonstrate that the TLR assay is able to distinguish between normal and abnormal DDR function, and we therefore proceeded by testing missense variants in PALB2.

### Validation of the TLR assay for clinical interpretation of *PALB2* missense variants

Previous studies in *PALB2* have primarily been domain-specific.^17,36,37^ In order to validate an assay for PALB2 as a whole, we selected variants across the length of the protein (Table S7). As controls, nine benign/likely benign missense variants (N241D, I309V, Q559R, E672Q, I676T, A712V, P864S, V932M, and G998E) were selected from the Clinical Variant Database (ClinVar)^9^ and manually reviewed to ensure that the interpretation could be reached confidently without using functional evidence. As there are currently no definitive likely pathogenic/pathogenic missense variants in PALB2, we selected three frameshift/early truncating variants (C77Vfs*100, N497Mfs*64, L531Cfs*30) that could reach at least a likely pathogenic classification without functional evidence to serve as controls. It should be noted that expression of these truncating variants from cDNA may not reflect the true biological context (in which nonsense mediated decay would be expected to occur), however these cDNA constructs should still produce a non-functioning protein given that a large portion of the C-terminal end of the protein will be absent. Additionally, there is evidence that the c.1592delT variant that encodes L531Cfs*30 escapes NMD but results in truncated protein that is highly unstable with low levels of protein product and functionally abnormal.^38^ In addition, 25 missense VUS were selected from ClinVar or the literature, or were synthetically designed. We examined the available evidence and prioritized those that did not have prior functional evidence (K18R, L24S, K30N, R37S, N186I, E331Q, F404L, P490A, A712P, N821Y, L939W, A1025T, I1037T, L1070P, P1097R), as these VUS would stand to benefit the most from the addition of new functional data. Other VUS selected from ClinVar include PALB2 variants Y28C, L35P, R37H, and T1030I. Synthetic variants and those selected from the literature include L17P, L21P^39^, L24P^39^, Y28P, R37P, and A1025R^40^. Typical flow plots for one run of the PALB2 adaptation of the TLR assay are represented in Figure 3.

**Figure 3.**
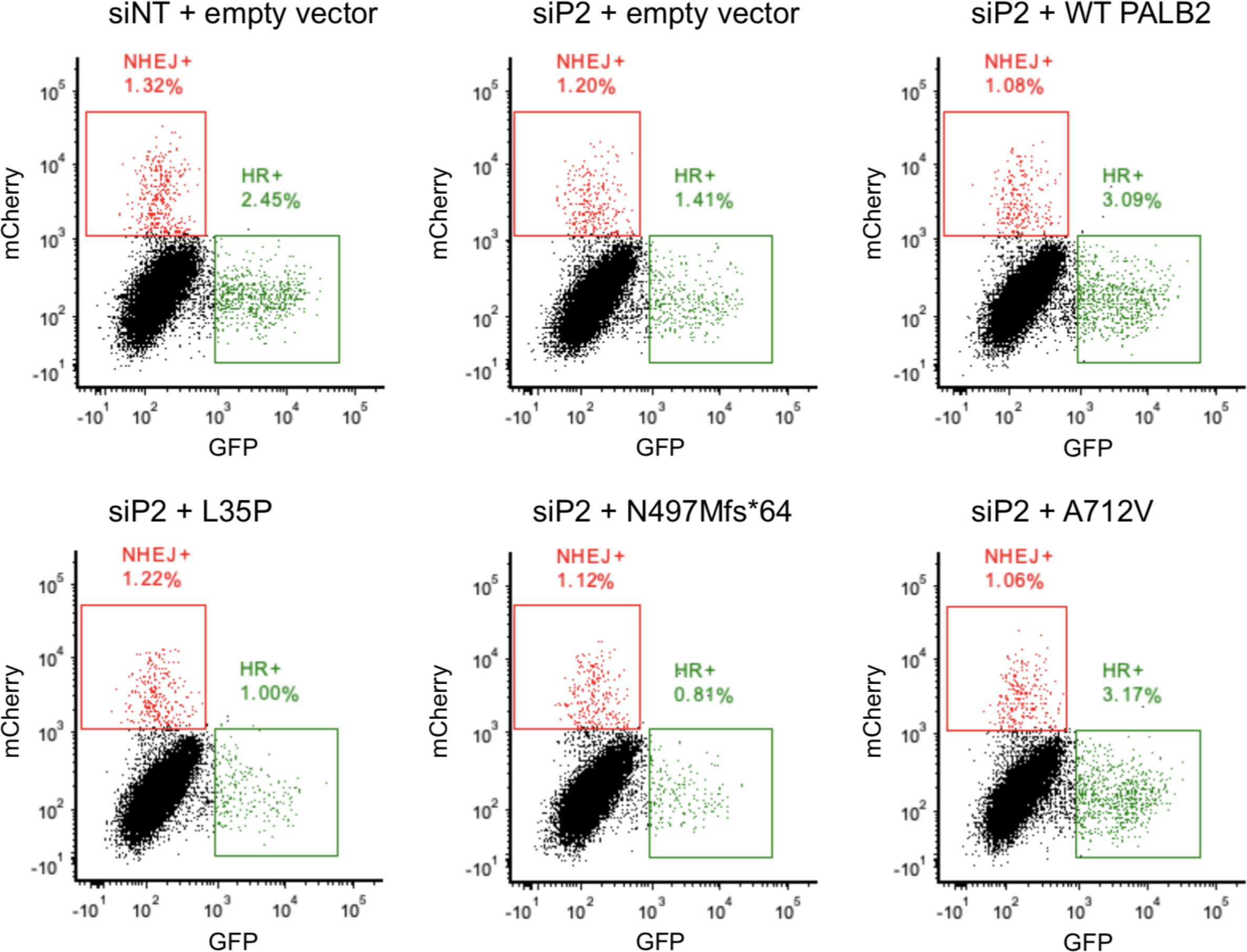
Representative PALB2 TLR assay flow cytometry data. Typical flow plots for one run of the TLR assay. **Row 1 (left to right):** Controls for siRNA/DNA transfection and batch effects. **Row 2 (left to right):** Transient rescue with PALB2 variants L35P (potentially pathogenic missense variant)^22^, N497Mfs*64 (known pathogenic), and A712V (known benign).

We followed the same experimental protocol as was developed for *BRCA2*, using PALB2 WT cDNA and EV as “normal” and “abnormal” controls, respectively. Each PALB2 variant was included in at least 3 independent experiments. We performed in-plate adjustment to the mock treated condition and LMM random effect normalization across all of the plates as described above. As shown in Figure 4a, we found that WT and benign PALB2 variants restore HR, indicated by a higher HR/NHEJ ratio (means range from 1.073 – 1.517) while EV and truncated PALB2 variants exhibit fewer HR events, with a lower HR/NHEJ ratio (means range from 0.285 – 0.721). In the PALB2 experiments, the GMM distribution that corresponds to WT controls had a mean HR/NHEJ of 1.249 (standard deviation 0.262) and the distribution that corresponds to the EV controls had a mean HR/NHEJ of 0.554 (standard deviation 0.098). Again, we applied thresholds based on a 10:1 ratio of the probability to belong to either the normal (HR/NHEJ > 0.848) or abnormal (HR/NHEJ < 0.689) distributions (Figure S3). Using these thresholds, the mean HR/NHEJ assay read-out of all PALB2 B/LB validation controls fell in the normal range; two of the truncating variants (N497Mfs*64 and L531Cfs*30) had mean HR/NHEJ assay values in the abnormal range, while one truncating variant (C77Vfs*100) was considered of intermediate or indeterminate functional impact. Therefore, the TLR correctly classified all nine PALB2 B/LB controls and two out of three P/LP controls, which corresponds to an OddsPath of 6 (PS3_moderate) for an abnormal functional readout and an OddsPath of 0.333 (BS3_supporting) for a normal functional readout. Assay results are depicted for variants across the entire PALB2 protein (Figure 4b).

**Figure 4.**
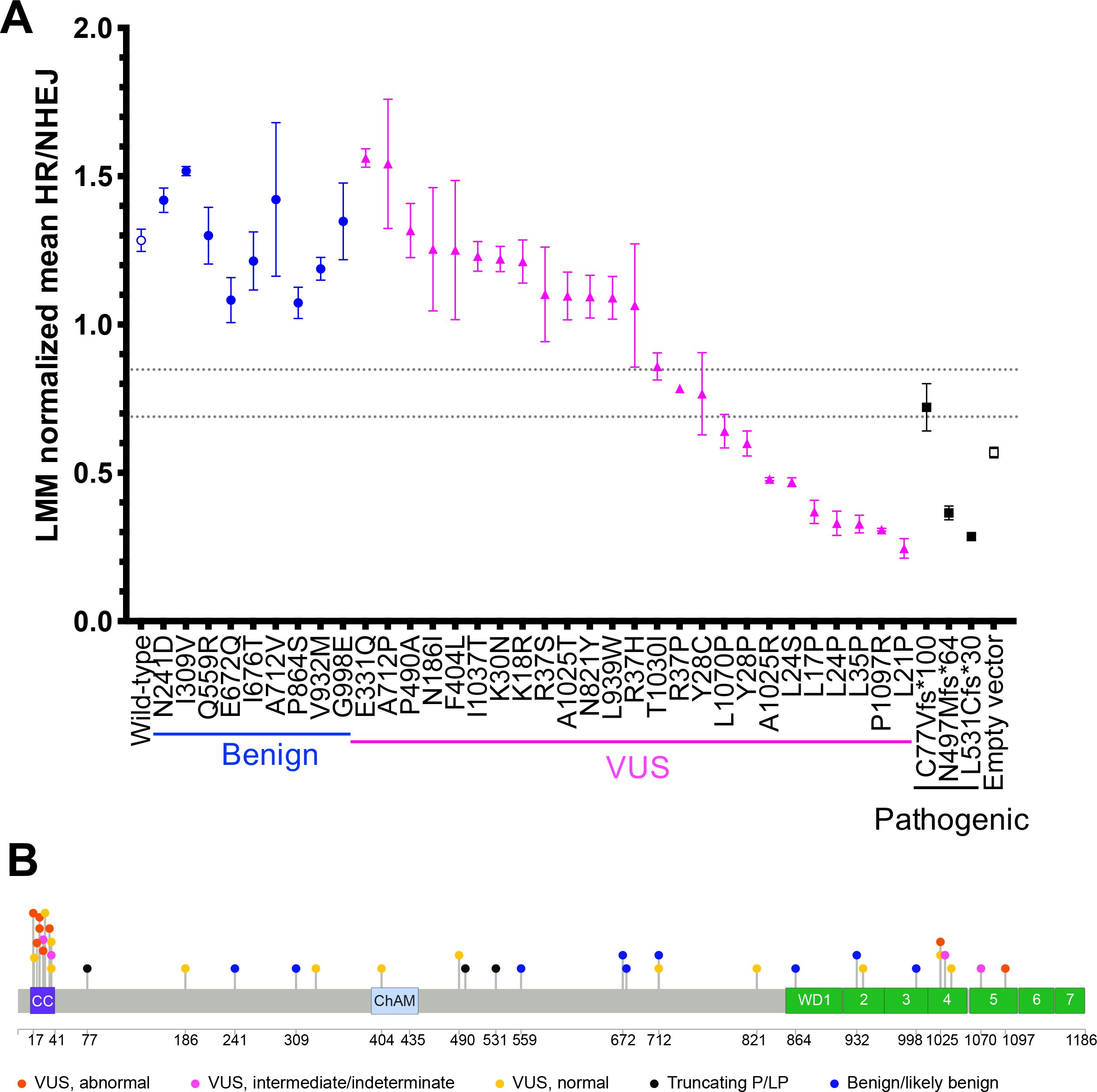
Summary of PALB2 TLR assay read-out for all variants tested. (A) TLR assay results for the PALB2 experiments are summarized, demonstrating HR function after siPALB2 knockdown and rescue with plasmids containing PALB2 variants or controls. HR function is rescued by PALB2 wild-type cDNA and benign validation controls but is impaired with empty vector and pathogenic validation controls. Each rescue condition is depicted across the x-axis. The y-axis represents the LMM normalized HR/NHEJ ratio as the mean ± SEM of at least three independent experiments. For wild-type and empty vector controls, N=40. Thresholds for normal and abnormal functional impact are indicated by gray dashed lines. (B) Lollipop diagram of PALB2 protein variants tested in the TLR assay, colored by functional impact. Benign validation controls are shown in blue, pathogenic validation controls are shown in black. VUS are coded by functional impact, based on the thresholds set for “normal” (yellow) or “abnormal” (red) assay read-out. VUS with intermediate assay read-out are labeled in pink. Coiled-coil (CC), chromatin association motif (ChAM), and WD40 protein domains (WD1-7, seven blades of the WD beta propeller) are indicated by colored boxes.

### Functional assessment of PALB2 missense VUS

PALB2 missense VUS span the range of functional readout from normal to abnormal, with some of intermediate or indeterminate function. Interestingly, we found substantial effects on DDR for eight VUS with mean HR/NHEJ ratios between 0.245 – 0.599. Consistent with Foo and colleagues’ original report^22^, as well as subsequent investigations^23–25^, we found that L35P had a robust abnormal functional impact on HR/NHEJ. Other variants displaying abnormal function include: L17P, L21P, L24P, L24S, Y28P, A1025R, and P1097R. We also found four PALB2 missense variants with intermediate function (Y28C, R37P, T1030I, and L1070P), with mean HR/NHEJ ratios between 0.641 – 0.859. While T1030I and L1070P could be assessed based on mean HR/NHEJ as having normal and abnormal function, respectively, the SEM for each extended into the indeterminate range. For this reason, we opted for the more conservative interpretation of “indeterminate” functional readout from the perspective of applying functional evidence for variant classification. It is likely that additional replicates for these variants would provide a more precise estimate of the mean HR/NHEJ value. The remaining 13 VUS tested performed above the threshold for normal function (mean HR/NHEJ: 1.064 – 1.562). Overall, these results indicate that the TLR assay is able to distinguish missense variants with normal and abnormal repair functions. Our results also suggest that some missense variation in the BRCA1 and BRCA2-interacting domains of PALB2 may have deleterious effects on DDR, as seen previously in truncating PALB2 variants.

Next, we performed follow-up evaluation of protein re-expression by western blotting for FLAG-tagged missense PALB2 variants, including seven with abnormal HR/NHEJ, as well as EV and a benign variant control, N241D. EV control does not express FLAG-tagged PALB2. These seven VUS had detectable expression, similar to N241D and WT PALB2 (Figure S1d). For L17P, L21P, L24P, L24S, Y28P, L35P, and P1097R, this suggests that their abnormal HR/NHEJ ratios are not due to absent expression or reduced protein stability. Consistent with our results, Boonen et al. recently demonstrated normal protein expression for L24S and L35P variants.^25^ They also found normal protein expression for K18R, Y28C, R37H, L531Cfs*30, Q559R, E672Q, P864S, L939W, G998E, and A1025R, but decreased expression/stability for T1030I, I1037T, and L1070P.^25^

### TLR readout is concordant with other assays of HR in PALB2

While this manuscript was in preparation, Boonen^25^, Wiltshire^23^, and colleagues published manuscripts that examined HR capacity of PALB2 variants using the DR-GFP fluorescent reporter system in *Trp53^KO^/Palb2^KO^* mouse cell lines, either mammary tumor or mouse embryonic stem cells (Table S7). We therefore asked how well the readout of our TLR assay correlates with the results for missense variants tested across these other two assays by linear regression (Figure 5). When plotted against TLR mean HR/NHEJ, R-squared values from these regressions (0.7220 and 0.7564) suggest that our results are generally consistent with the published DR-GFP data (Figure 5, top row). We also considered that using the TLR readout of %HR might make a more direct comparison, and repeated the regressions with this data. Interestingly, the R^2^ value decreased slightly to 0.6544 and 0.7207 for the Boonen et al.^25^ and Wiltshire et al.^23^ datasets, respectively (Figure 5, bottom row). Overall, the positive correlation of TLR HR/NHEJ readout with other data in the literature increases our confidence in this metric of functional impact for PALB2 missense variants.

**Figure 5.**
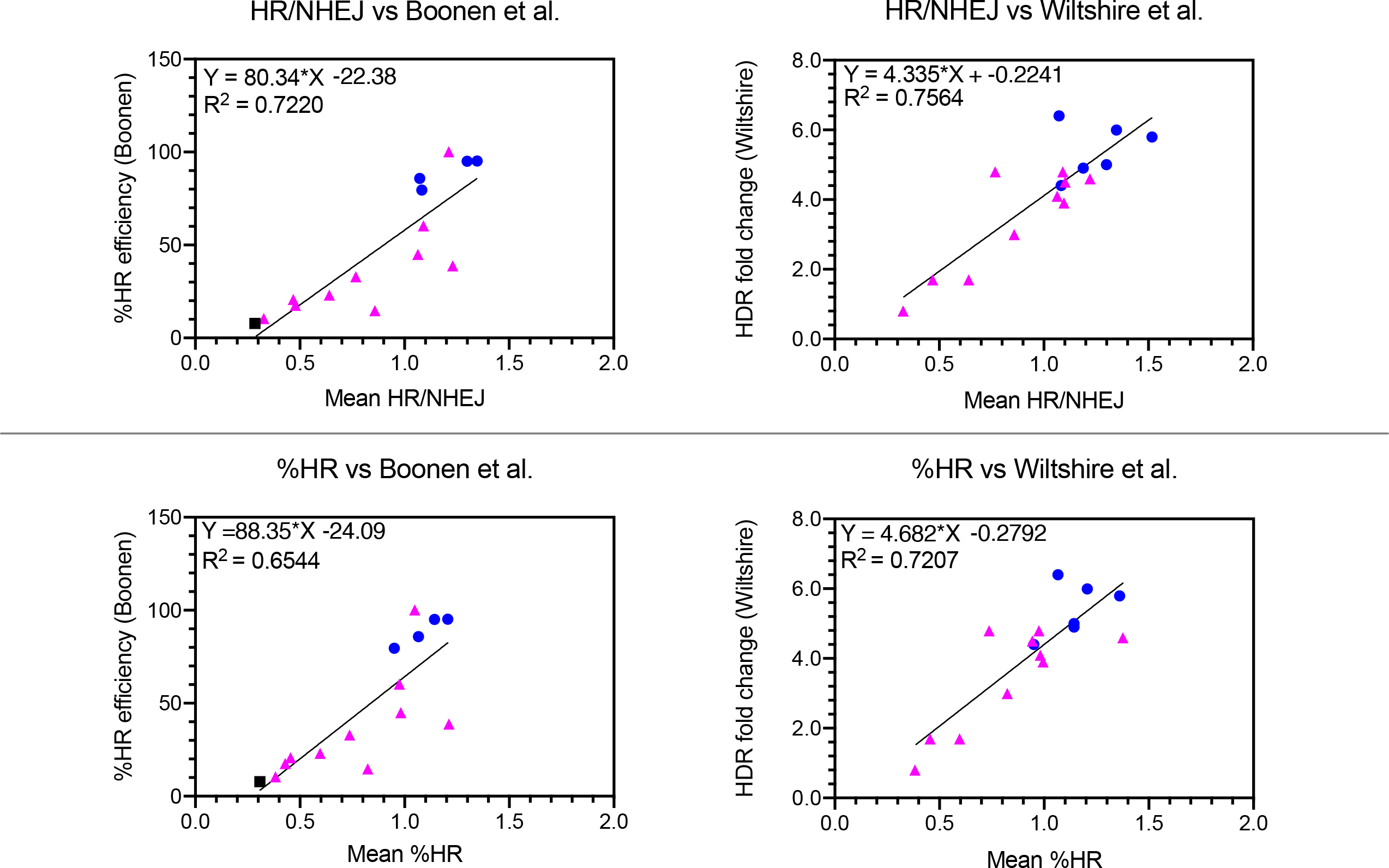
TLR assay HR/NHEJ values correlate with published HR functional data. Top row (left to right): Linear regression of normalized mean HR/NHEJ versus %HR efficiency (Boonen et al.^25^, N=15), or HDR fold change (Wiltshire et al.^23^, N=16). Bottom row (left to right): Linear regression as above, but using normalized mean %HR instead of HR/NHEJ ratio. Benign/likely benign variant controls are shown as blue circles, VUS as magenta triangles, and truncating variants as black squares.

## DISCUSSION

To assess the functional impact of missense variants in *PALB2* for clinical variant interpretation, we developed and validated the TLR cellular *in vivo* fluorescent reporter assay of HR and alt-NHEJ DDR outcomes. We first determined the TLR’s performance using 16 BRCA2 missense variants as benchmarks for normal/abnormal HR function. Our results for these controls were consistent with previous studies.^27,41,42^ We then tested 37 PALB2 variants, generating functional data for germline *PALB2* variants at a moderate level of evidence for a pathogenic interpretation (PS3_moderate) or a supporting level of evidence in favor of a benign interpretation (BS3_supporting), based on the ability of the assay to correctly classify 9 B/LB validation control variants and 2 out of 3 P/LP validation control variants.

PALB2 missense VUS seem to cluster by normal and abnormal functional impact, with a few having intermediate or indeterminate function. Of the 25 PALB2 missense VUS examined, eight demonstrated abnormal function comparable to EV and truncating variants. These variants are all located in either the N-terminal coiled-coil (CC) domain or the C-terminal WD40 domain of PALB2, suggesting that these regions are critical for normal PALB2 DDR activity and may be hot spots for pathogenic missense variation.

### Coiled-coil domain and BRCA1 interaction

The CC domain (residues 9-44) is made up of a heptad of supercoiled alpha helices and is the site of PALB2 homodimerization, as well as heterodimerization with BRCA1.^10,43^ Alpha helical structures are disrupted by proline substitutions.^15^ As such, it is unsurprising that most of the CC domain variants where the amino acid changed to a proline (L17P, L21P, L24P, Y28P, L35P) had an abnormal readout. Variants Y28C and R37P were of intermediate functional effect. Based on previous work, L17, L21, L24, Y28, T31 and L35 are the key residues of the PALB2-PALB2 interface.^15^ The exact role of PALB2 homodimerization in regulating HR is still unclear, as L17, L21, L24, Y28, and L35 also make up the hydrophobic interface with BRCA1. Recently, two other groups found L24S to have abnormal HR function, in addition to decreased RAD51 foci, some increased sensitivity to poly(ADP-ribose) polymerase inhibitors (PARPi), and impaired interaction/complexing with BRCA1.^23,25^ They also found L24S to have normal protein expression, consistent with our findings (Figure S1d), normal nuclear localization, and even increased protein stability in one report.^23,25^ This suggests that defects in DNA damage repair are likely mediated by impaired BRCA1 interaction, rather than issues of protein expression, stability, or localization.

PALB2 variant Y28C fell in the range of intermediate functional impact (mean HR/NHEJ of 0.767) in the TLR assay. Data in the literature for Y28C have not reached consensus on its functional impact. Wiltshire et al. considered it to have normal HR activity (fold change of 4.8) in a mES cell DR-GFP assay^23^, but Boonen et al. considered Y28C to have intermediate function (32.92% HR efficiency) in the same assay^25^. Rodrigue and colleagues showed between 25-40% residual HR activity by the CRISPR-LMNA assay^24^, while Foo et al. found a 65% reduction in HR activity (35% of normal) and decreased PALB2-BRCA1 co-immunoprecipitation (co-IP), but normal recruitment to BRCA1 foci.^22^

PALB2 residues K18, K30, and R37 are also part of the coiled-coil protein-interacting domain, but are not points of direct contact.^22^ This may help explain why certain CC domain residues we tested may be more sensitive to alterations than others. K18R is listed as having conflicting interpretations in ClinVar (Accession: VCV000126758.3), with a combination of likely benign, benign, and VUS classifications reported from different clinical laboratories. In addition to its normal HR function demonstrated here and elsewhere^23–25^, its MAF is too high to be considered pathogenic for hereditary breast cancer, with a filtering allele frequency of 0.01733 in gnomAD v2.1.1 (accessed 7/4/2020)^44^, suggesting that this variant should be considered benign (BA1, BS3_supporting). While K30N has been reported in 1/747 Australasian females with multiple-case breast cancer families^45^, we and others found it to have normal HR function^22,23^ although the level of evidence that is currently warranted by our assay (BS3_supporting) is not sufficient to cause it to be reclassified.

Variants R37H and R37S exhibited normal function based on the thresholds we applied. Available functional literature for R37H is generally consistent with our interpretation^22,23,25^, though one group reported ~40% of normal HR, congruent with a mild increase in olaparib sensitivity^24^. We have previously observed R37S in exomes of two research participants evaluated for non-cancer-related indications (cardiomyopathy; neurodevelopmental disorder with dysmorphic features). In the literature, R37S has been reported in a patient with breast cancer and it was noted that it may have an effect on gene splicing^46^, though according to one report in ClinVar, “multiple internal splicing models do not predict an effect on splicing” (Accession: SCV000565339.3). For variants where we have demonstrated a supporting level of evidence for normal DDR activity, it is important to note that we have not assessed all functions of PALB2 that may lead to disease pathogenesis, such as splicing, and thus caution is still merited in applying benign functional criteria for clinical variant interpretation. R37P (mean HR/NHEJ of 0.785) was considered of indeterminate function by the TLR assay. Given what is known about the impact of prolines in the CC domain, it is not surprising that R37P appeared to perform at a slightly lower level than other amino acid substitutions at the same residue, though it is unclear if this is sufficient to have a meaningful clinical impact at the R37 residue.

Interestingly, pathogenic frameshift variant C77Vfs*100 appeared to have a higher mean HR/NHEJ ratio (0.721) than EV alone (0.568), falling in the intermediate range. Tischkowitz and colleagues first described this frameshift variant in a family of Scottish ancestry with strong history of breast cancer—7 breast cancers in 3 female mutation carriers. They found c.229delT (p.C77Vfs*100) had decreased HR activity and increased sensitivity to mitomycin C (MMC), a DNA cross-linking agent.^47^ This alteration is predicted to result in loss of normal protein function through either protein truncation or nonsense mediated decay, as the frameshift leads to the creation of a premature stop codon at position 100 in the new reading frame. The truncated protein includes the complete CC domain of PALB2, which would permit interaction with BRCA1 and PALB2-PALB2 binding. While this is also true of the other frameshift variants examined, it is possible that our artificial system led to overexpression of an artificial protein fragment that retained enough of the C-terminal function to affect the relative HR/NHEJ. Alternatively, our result may reflect the true range of HR/NHEJ activity for pathogenic variants.

### WD40 domain and BRCA2 interaction

Two variants in the PALB2 C-terminal WD40 domain, A1025R and P1097R, demonstrated robust abnormal HR/NHEJ functional read-out in the TLR assay. The WD40 domain is a ring-like beta-propeller structure common in protein-protein interactions^40^. PALB2 residues V1019, M1022, A1025, I1037, L1046, K1047, L1070, P1097, and K1098 line a hydrophobic pocket of the WD40 domain that binds BRCA2^40^; residues A1025 and P1097 are located in the 4^th^ and 5^th^ blades^36,40^, respectively, suggesting that the observed abnormalities in HR activity may be related to impaired BRCA2 interaction. Truncation of PALB2 after P1097 abrogates BRCA2 binding^36^, further implicating this domain in BRCA2 interactions and HR activity.

Interestingly, prediction algorithms for the impact on splicing suggest that P1097R may activate a cryptic exonic splice donor site.^48^ Although our results suggest that the missense change itself results in abnormal HR function, splicing abnormalities cannot be assessed using the cDNA constructs used here, so we cannot rule out an alternative splice-disrupting mechanism that may occur in the context of the endogenous gene.

We also observed abnormal HR/NHEJ function for A1025R, consistent with previous reports.^25,49^ It has also been shown that A1025R abrogates the PALB2-BRCA2 interaction^23,24,37,40,49,50^, increases sensitivity to olaparib and cisplatin^23,25^, and affects progression through the G2/M checkpoint^25,51^. However, A1025R does not seem to impact PALB2 protein stability^25,40^. Of note, a change to threonine at the same A1025 residue had normal readout in our assay and in the DR-GFP assay in mouse cell lines^23^. Threonine is a relatively conservative physiochemical change and multiple mammalian species have threonine at this position (ClinVar Accession: SCV000290863.4), suggesting that it does not substantially alter protein function. Interestingly, the only two PALB2 variants in the WD40 domain with clear abnormal functional consequences (A1025R and P1097R) were alterations predicted to substitute an arginine for the native residue. Future studies to elucidate which amino acid substitutions are tolerated in this region would be pertinent.

I1037T also affects a residue located in the hydrophobic pocket of PALB2 that binds BRCA2^40^, but demonstrated inconsistent results between different assays. We found that I1037T did not have a marked effect on HR/NHEJ in the TLR assay, whereas an intermediate impact on HR (38.86% efficiency) was observed by DR-GFP in *Trp53^KO^, Palb2^KO^* mES cells.^25^ In the same study, I1037T showed decreased protein expression, but normal mRNA expression and normal PARPi sensitivity.^25^ The incongruency between the TLR and DR-GFP assays may reflect differences in thresholding that could be resolved by benchmarking the assay readout with variant controls.

L1070P is also located in the hydrophobic pocket and was just below the threshold for abnormal functional impact with a mean HR/NHEJ of 0.641, though its SEM crossed into the indeterminate range. Other studies considered L1070P to have defects in HR, with 23.09% HR efficiency in one study^25^ and an HDR fold change of 1.7 (equivalent to less than 34% of normal) in another.^23^ Recent work has further shown L1070P to impact protein expression/stability, potentially increase sensitivity to olaparib and cisplatin, partially disrupt the BRCA2 interaction, decrease PALB2 recruitment to DSBs, and increase cytoplasmic retention.^23,25^ Another PALB2 variant in the WD40 region, T1030I, also demonstrated an intermediate, bordering on normal, functional impact on HR/NHEJ (0.859) in the TLR assay. Wiltshire’s DR-GFP assay (fold change of 3.0)^23^ agreed with this intermediate classification, though Rodrigue’s CRISPR-LMNA assay^24^ and Boonen et al.’s DR-GFP assay^25^ considered it to have abnormal function (23.6% and 14.68% of normal HR, respectively). T1030I has demonstrated modest impacts on RAD51 and RAD51C interactions^24,36^ and PARPi sensitivity^24,25^ and no defect in BRCA2 complex formation.^24^ T1030 is not one of the key residues in the hydrophobic pocket that interacts with BRCA2, and others have found no defect in BRCA2 complex formation. There are reports that T1030I has decreased protein stability^25,36^ and one group found increased localization to the cytoplasm^24^, which could help explain some decreased HR function, depending on how much T1030I is active in the nucleus. The overall functional impact of T1030I remains unclear.

Additional replication of the results for L1070P and T1030I in the TLR assay may further improve our estimate of their mean HR/NHEJ read-out; alternatively, these conflicting results may indicate a limitation of the dynamic range of the current TLR assay to discriminate between normal and abnormal functional impact for all variants. It is also possible that T1030I, L1070P, and other variants of intermediate function are hypomorphic, which could be interpreted as having a modestly increased risk of cancer, or an increased risk of a less severe cancer phenotype. There is some precedent for hypomorphic variants in *BRCA2^52^* conferring modest increases in cancer risk. There are also hypomorphic *PALB2* variants that have been associated with a less severe Fanconi Anemia (MIM: 610832) phenotype.^53^ Family studies and correlation with clinical phenotype could help resolve these questions moving forward.

A final PALB2 VUS in the WD40 domain, L939W, revealed normal HR/NHEJ function in the TLR assay. L939W is listed as having conflicting interpretations in ClinVar, with a combination of likely benign, benign, and VUS classifications reported from different clinical laboratories. There are previous reports that L939W displays a modest decrease in BRCA2 and RAD51 binding, and affects HR capacity^36^, but others have been unable to reproduce this HR finding^23,25,54^. Given that the original studies of abnormal activity were conducted without validation controls, these results do not qualify as strong PS3 evidence and should be interpreted cautiously in light of substantial refuting evidence. Located in the second blade of the WD40, L939 is not in direct interaction with BRCA2. We have observed L939W in four exomes analyzed for non-cancer-related indications, including cardiomyopathy, neuromuscular disorders, dysmorphology, and neurodevelopmental disorders. Furthermore, the reported filtering allele frequency of 0.001421 in gnomAD (accessed 7/4/2020) is too high to be considered pathogenic for hereditary breast cancer.^44^ We therefore support a likely benign classification for this variant (BS1, BS3_supporting).

### Rigorous assay thresholding for result concordance

While the majority of our results are consistent with published studies (Figure 5), most discrepant interpretations involve variants that are of intermediate function. The overall interpretation depends on each assay’s thresholds for calling the readout “abnormal”, “normal”, or “intermediate”. Comparing results is further complicated if some assays use a binary classifier, eliminating an “intermediate function” category. We also note that while other assays normalize their functional read-out to the WT control, this approach masks the true variation in WT readout across separate experimental replicates and creates potential challenges when drawing thresholds for normal versus abnormal function. In our assay, we instead used WT and EV controls to set thresholds for assay read-out ranges that could be considered normal, intermediate, or abnormal based on a GMM. This allowed us to then benchmark our assay based on the results for several validation controls, consistent with guidance for validating functional assays and setting strength thresholds for evidence application developed in collaboration with the ClinGen Sequence Variant Interpretation Working Group^8^. The discrepancies in the interpretation of DDR assay readout highlight the importance of rigorous assay design to enable reproducible statistical thresholding.

### Conclusions

The work presented here represents the first validated functional assay for *PALB2* with statistically determined thresholds of evidence strength for application of PS3/BS3 criteria in clinical variant interpretation. Our novel approach to validation using 16 known benign and pathogenic variants in *BRCA2* to establish the protocol for this assay demonstrates that pathway-based approaches for assay development are promising when benchmarking controls are limited. We also used nine B/LB PALB2 missense controls and 3 truncating P/LP variants to further benchmark the TLR assay for PALB2. While results for variants with normal readout can be used at a supporting level of evidence in favor of a benign classification (BS3_supporting) and abnormal readouts as moderate evidence supporting pathogenicity (PS3_moderate), including additional variants of known significance (as they become available) would further increase the strength of evidence that can be applied. This limitation is unfortunately an inherent problem for all assays of PALB2 function, given the paucity of pathogenic missense variants that can be used for clinical validation.

It is now becoming clear that L35P is not the only PALB2 missense variant with abnormal HR function – others in the BRCA1- and BRCA2-interacting domains have now demonstrated abnormal functional impact on DDR outcomes in multiple different assays. As such, further exploration of missense variation in these domains is merited. However, additional clinical observations and family segregation evidence, in addition to this moderate strength functional evidence, will be needed in order to conclude that these variants are pathogenic. Adapting the TLR assay for multiplexed assessment of *PALB2* VUS would permit evaluation of larger numbers of variants before they are even seen in a patient, preempting delays in conclusive variant interpretation while functional evidence is generated. Overall, we expect that the new functional data presented here will contribute the additional evidence required to help reclassify these VUS within the ACMG/AMP variant interpretation framework for direct clinical benefit.

## Supplemental Data

Supplemental Data include Supplemental Material and Methods, three figures, and seven tables.

## Acknowledgments

The authors would like to thank Stephanie Bellendir Crowley, PhD, for her assistance in molecular cloning and preliminary validation experiments.

JSB is a recipient of the Yang Family Biomedical Scholars Award. SEB was supported in part by the National Institute of General Medical Sciences grants 5T32 GM007092 and 5T32 GM008719-6. ECA received support in part from the Chancellor’s Science Scholars fund. The UNC Flow Cytometry Core Facility is supported in part by P30 CA016086 Cancer Center Core Support Grant to the UNC Lineberger Comprehensive Cancer Center, as well as Center for AIDS Research award number 5P30AI050410.

## Web Resources

ClinVar, https://www.ncbi.nlm.nih.gov/clinvar/

gnomAD, https://gnomad.broadinstitute.org/

OMIM, https://www.omim.org

## Data Availability

Data supporting the findings presented here are available from the author upon reasonable request.

