## Supplemental Methods and Data for "A Validated Functional Analysis of PALB2 Missense Variants for Use in Clinical Variant Interpretation"

### SUPPLEMENTAL MATERIALS AND METHODS

#### TLR Assay Optimization

#### DSB repair timing

In order to determine the optimal timing for DNA transfection and flow cytometry analysis, 5.0 x $10^5$ cells were reverse transfected in a 12-well plate with 0.5 μg of I-SceI-IFP and 0.5 μg of GFPdonor-BFP using Lipofectamine 2000 (Invitrogen) at 12, 24, 48, or 72 hours before flow cytometry. Controls included untreated cells and cells transfected with 0.5 μg of empty vector only. To track DNA transfection efficiency and timing of plasmid expression, $IFP^+$/$BFP^+$ cells were analyzed as a percentage of single, live cells. DSB repair timing was measured by the percent of cells that were $GFP^+$ (%HR) or $mCherry^+$ (%NHEJ). At least 100,000 single, live cells were analyzed in duplicate for each condition (Figure S1a).

#### Timing siRNA-mediated knockdown

We used siRNA against the human ubiquitin B (*UBB*) gene (MIM: 191339), which results in cell death upon knockdown, to verify siRNA knockdown during assay development (M-013382-01-005, Dharmacon). As a control in all knockdown experiments, we used a non-targeting siRNA (siNT) pool (D-001206-13-05, Dharmacon). Individual siRNAs were used for *PALB2*[1] and *BRCA2* silencing (D-012928-04 and D-003462-02, respectively, Dharmacon) (Table S2).

In order to determine the timing of siRNA-mediated knockdown, 3.0 x $10^5$ 293T/TLR cells were reverse transfected with 75 nM of siRNA (either siNT, siPALB2, or siUBB) per well of a 6-well plate using Lipofectamine RNAiMAX (Invitrogen) according to the manufacturer's protocol. Cells were harvested by enzymatic release after 24, 48, 72, or 96 hours. Pellets were stored at -20°C until RNA or protein extraction was performed. mRNA expression was analyzed by RT-PCR for three independent experiments (Figure S1b). For protein analysis, western blots were performed using 15 μg of whole-cell lysate per well, with siNT-treated cells serving as a control for each timepoint (Figure S1c).

**Assessing transient re-expression of PALB2 variants**

To evaluate transient PALB2 re-expression, 3.0 x $10^5$ 293T/TLR cells were treated with 75 nM siPALB2, followed by co-transfection 24-hours later with 1.25 μg of I-SceI, 1.25 μg of GFPdonor, and 0.71 μg of the indicated pDEST-FRT/T0-FLAG-PALB2 construct. Cells were harvested by enzymatic release 48 hours after the siRNA transfection and pellets were stored at -20°C until protein extraction. For protein analysis, western blots probing for FLAG-tagged PALB2 were performed using 25 μg of whole-cell lysate per run. WT PALB2 served as a control on each blot (Figure S1c).

### SUPPLEMENTAL FIGURES

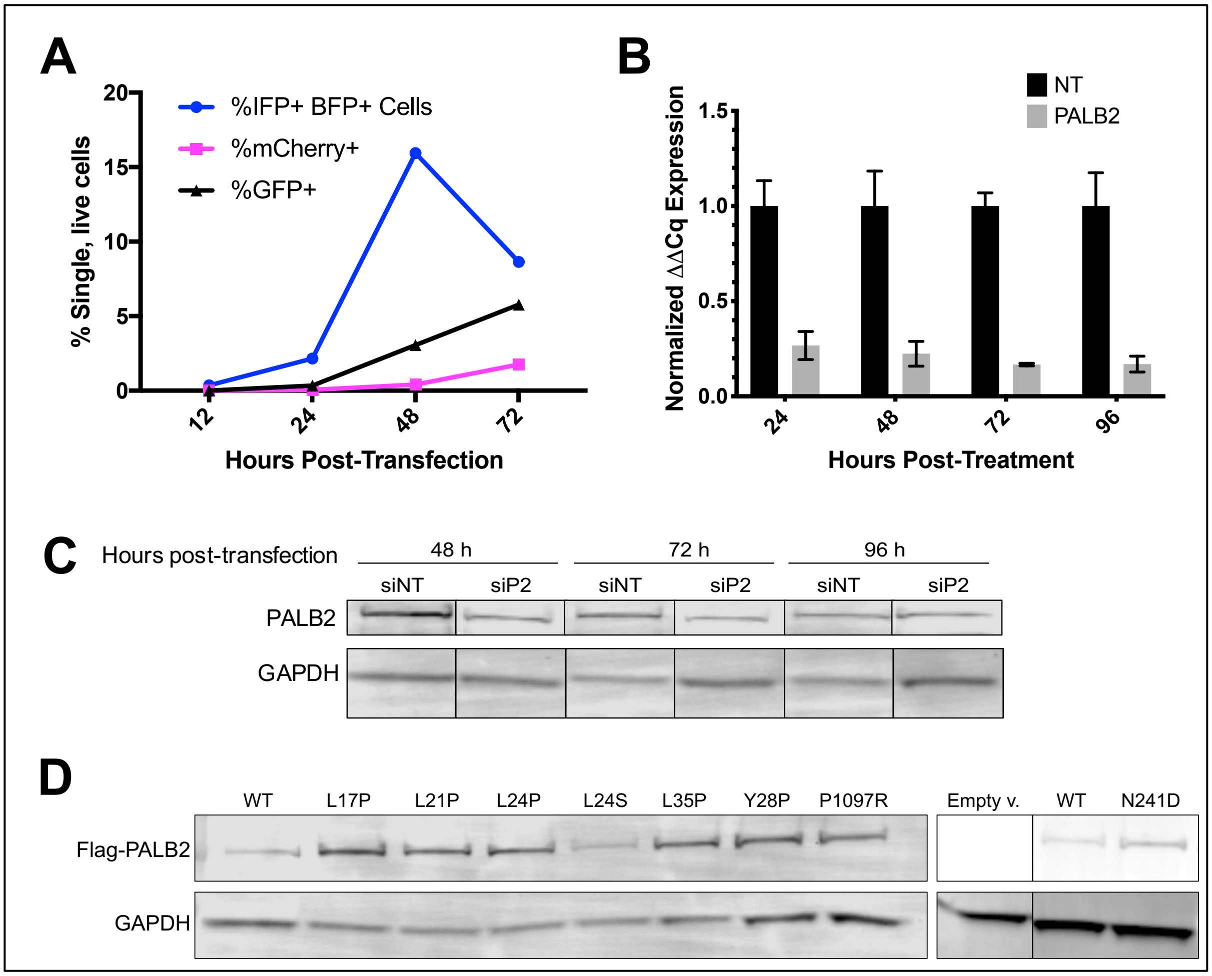


**Figure S1: Development of TLR assay protocol.**
(A) Timing DSB repair by flow cytometry. 293T/TLR cells transfected with I-SceI-IFP and truncGFP Donor-BFP at 12, 24, 48, or 72 hours before flow cytometry was performed. $IFP^+/BFP^+$ cells (doubly transfected with both nuclease and truncGFP), are shown as percent of single, live cells. mCherry and GFP expression serve as markers for alt-NHEJ and HR events, respectively; (B) Time course of siRNA-mediated knockdown of *PALB2*. Expression of *PALB2* mRNA after siRNA-mediated knockdown relative to non-targeting control siRNA (siNT) treated cells. 293T/TLR cells were transfected with 75 nM siRNA and cells were pelleted after the indicated number of hours and frozen for later RNA extraction and analysis. RT-PCR was performed using primers for *PALB2* and *B2M*. Results from 3 independent experiments shown as mean with standard deviation; (C) Western blot analysis of whole-cell lysates from 293T/TLR cells 48-, 72-, or 96-hours (h) after transfection with siNT or siRNA against *PALB2* (siPALB2). 15 μg of whole-cell lysate was loaded per lane and endogenous PALB2 expression was probed with 0.5 μg/ml anti-PALB2 antibody. GAPDH serves as a loading control. Lanes reordered here for ease of comparison against siNT control; (D) PALB2 protein re-expression. Western blot analysis of whole-cell lysates from 293T/TLR cells treated with 75 nM siPALB2 and transiently transfected 24-hours later with empty vector (empty v.) or pDEST-FRT/T0-FLAG-PALB2 encoding wild-type (WT) or variant PALB2. 48-hours after siRNA transfection, exogenous PALB2 expression was probed with 1.0 μg/ml anti-FLAG antibody. GAPDH serves as a loading control.

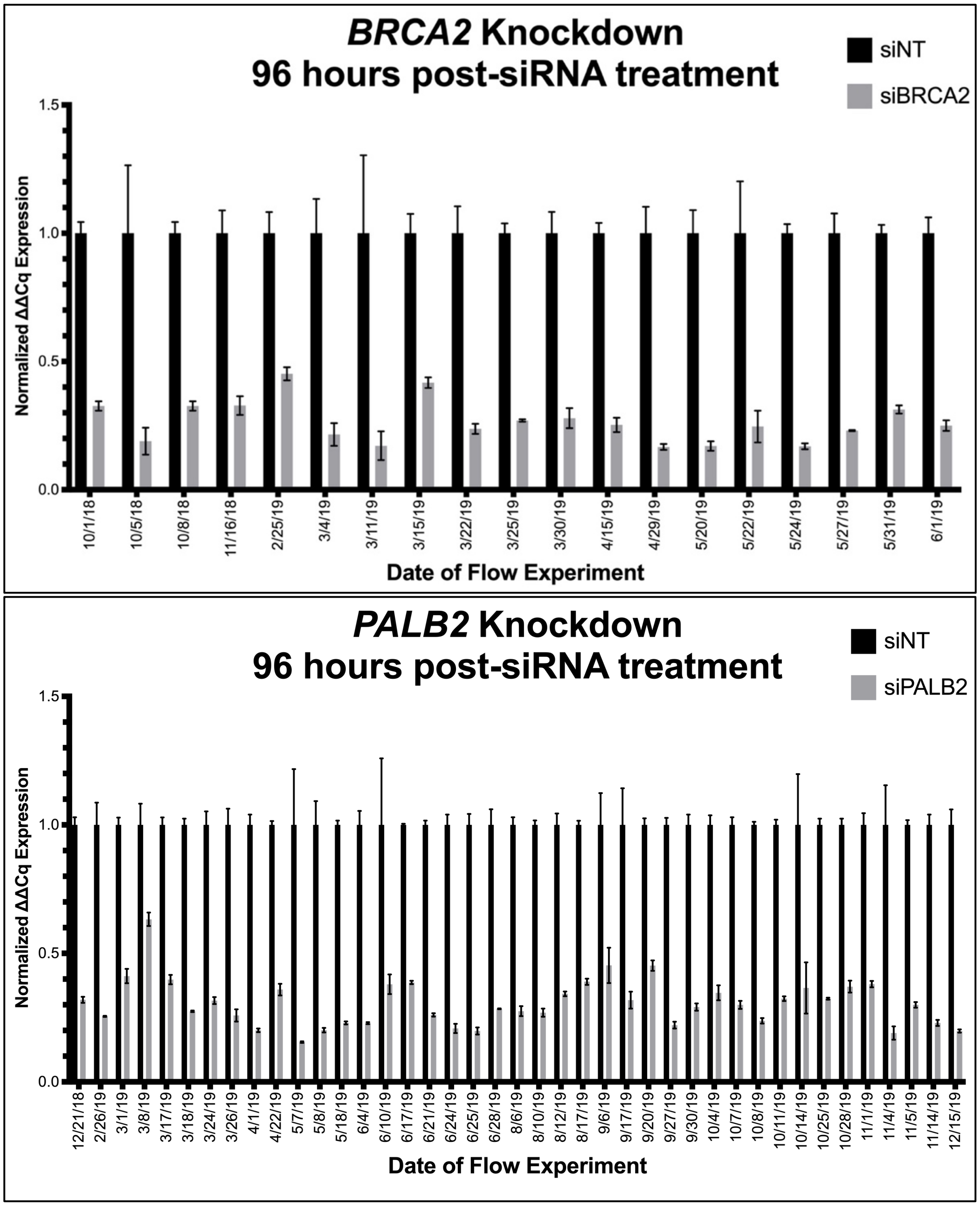


**Figure S2: Monitoring siRNA knockdown in flow cytometry experiments**. Results of RT-PCR performed in triplicate on 293T/TLR cells treated with either siNT, siBRCA2, or siPALB2 and rescued with empty vector and pelleted at the time of flow cytometry, 96 hours after siRNA transfection. Probes used: *B2M*, *BRCA2*, *PALB2*. All experiments demonstrated >50% knockdown of the target gene relative to siNT-treated cells at the time of flow. The only exception was the 3/8/2019 PALB2 flow sample, which appears to have had poor RNA integrity, as this RT-PCR result does not reflect the level of knockdown observed by flow cytometry (LMM normalized HR/NHEJ of 0.569 for empty vector rescue).

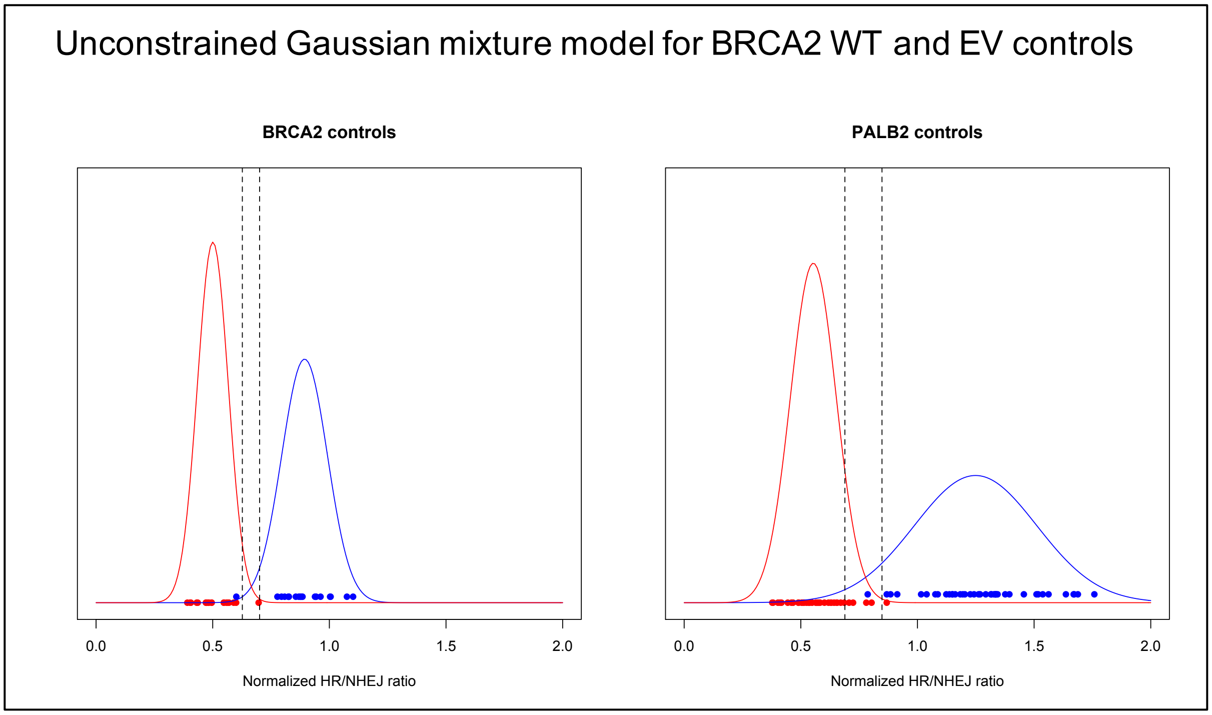


**Figure S3: Gaussian mixture models**

Each circle represents the TLR assay read-out for wild type (blue) or empty vector (red) controls, with corresponding curves based on an unconstrained Gaussian mixture model that provides distributions for “normal” and “abnormal” assay read-out. Dashed lines indicate read-out values where there is a 10:1 ratio of the probability to belong to one curve versus the other.

| Gateway Primer | Primer Sequence |
|---|---|
| Forward | ggggacaagtttgtacaaaaaagcaggctccaccatgcctattggatccaaa |
| Reverse | ggggaccactttgtacaagaaagctgggtattagatatattttttagt |

**Table S1: Gateway cloning primers.** Oligonucleotide sequences used to introduce *att*B sites by touchdown, gradient polymerase chain reaction (PCR) for Gateway cloning of wild-type *BRCA2*.

| siRNA | Target Sequences |
|---|---|
| siUBB control pool | GCCGUACUCUUUCUGACUA, GUAUGCAGAUCUUCGUGAA, GACCAUCACUCUGGAGGUG, CCCAGUGACACCAUCGAAA |
| Non-targeting control pool | UAGCGACUAAACACAUCAA, UAAGGCUAUGAAGAGAUAC, AUGUAUUGGCCUGUAUUAG, AUGAACGUGAAUUGCUCAA |
| siBRCA2 | GUAAAGAAAUGCAGAAUUC |
| siPALB2 | GAAGUCACCUCACACAAAU |

**Table S2: siRNA target sequences.** siRNA oligonucleotides used for transient knockdown of human *UBB*, *BRCA2*, or *PALB2* gene expression, or as non-targeting control.

| Gene | Nucleotide change | Forward Mutagenesis Primer | Reverse Mutagenesis Primer |
|---|---|---|---|
| *BRCA2* | c.4773T>C, c.4779A>G, c.4785G>A | gctgccccaaagtgcaaagagatgcaaaattctctcaataatg | cattattgagagaattttgcatctctttgcactttggggcag |
| *PALB2* | c.1635A>G, c.1641C>A, c.1647C>T | ggtccaaggaagaggtcacatcacataaatatcagcacg | cgtgctgatatttatgtgatgtgacctcttccttggacc |

**Table S3: Site-directed mutagenesis primers for conferring siRNA-resistance.** Mutagenesis primers used to confer plasmid siRNA resistance. Human Genome Variant Nomenclature (HGVS) nucleotide numbering shown for *BRCA2* and *PALB2* reference sequences (GenBank: NM_000059.3 and NM_024675.3, respectively).

| Gene | Primer name | Sequence | Corresponding nucleotides (CCDS) |
|---|---|---|---|
| *BRCA2* | BRCA2_1F | atgcctattggatccaaag | 1-19 |
| *BRCA2* | BRCA2_1342R | gtggcaaagaattctctg | 1342-1325 |
| *BRCA2* | BRCA2_1207F | ctaaatggagcccagatg | 1207-1224 |
| *BRCA2* | BRCA2_2541R | tcttgaaggtgatgctac | 2541-2524 |
| *BRCA2* | BRCA2_2406F | ttatgaatctgatgttga | 2406-2423 |
| *BRCA2* | BRCA2_3748R | cctcactaatattctcaa | 3748-3731 |
| *BRCA2* | BRCA2_3610F | gcttctggttatttaaca | 3610-3627 |
| *BRCA2* | BRCA2_5059R | gtaatgaagtctgactcacag | 5059-5039 |
| *BRCA2* | BRCA2_4831F | gtgccacctaagctctta | 4831-4848 |
| *BRCA2* | BRCA2_6193R | gctttccacttgctgtac | 6193-6176 |
| *BRCA2* | BRCA2_6058F | gaacattcagaccagctc | 6058-6075 |
| *BRCA2* | BRCA2_7400R | gctactgcttgattggag | 7400-7383 |
| *BRCA2* | BRCA2_7251F | cagagttgaacagtgtgt | 7251-7268 |
| *BRCA2* | BRCA2_8580R | cttttgttgggcctccac | 8580-8563 |
| *BRCA2* | BRCA2_8434F | ggaggaaatgttggttgt | 8434-8451 |
| *BRCA2* | BRCA2_9769R | tctcccctttacaagact | 9769-9752 |
| *BRCA2* | BRCA2_9634F | ggaaacaagcttctgatg | 9634-9651 |
| *BRCA2* | BRCA2_10257R | ttagatatatttttttagttg | 10257-10238 |
| *PALB2* | P2_1F | atggacgagcctcccggg | 0-18 |
| *PALB2* | P2_1305R | gacatccaaatgactctg | 1305-1288 |
| *PALB2* | P2_1147F | ctggaagcaacctctcct | 1147-1164 |
| *PALB2* | P2_2505R | ggaatgtttatgcagctc | 2505-2488 |
| *PALB2* | P2_2370F | agtgtcaggcaggcaagg | 2370-2387 |
| *PALB2* | P2_3561R | ttatgaatagtggtatac | 3561-3544 |

**Table S4: Sanger sequencing primers**. Sanger sequencing primers used to confirm *BRCA2* and *PALB2* construct and variant sequences. Primer name ends with “F” for “forward” or “R” for “reverse” primer direction. Reference sequences used were: CCDS9344.1 (GenBank: NM_000059.3) for *BRCA2*, CCDS32406.1 (GenBank: NM_024675.3) for *PALB2*.

| Nucleotide change | Amino acid change | Forward Mutagenesis Primer | Reverse Mutagenesis Primer |
|---|---|---|---|
| c.3055C>G | p.(L1019V) | cagcttcaaataaggaaatcaaggtctctgaacataacattaagaag | cttcttaatgttatgttcagagaccttgatttccttatttgaagctg |
| c.4570T>G | p.(F1524V) | aacctactctgttgggtgttcatacagctagcggg | cccgctagctgtatgaacacccaacagagtaggtt |
| c.6220C>A | p.(H2074N) | agcaagtttccattttagaaagttccttaaacaaagttaagggagtg | cactcccttaactttgtttaaggaactttctaaaatggaaacttgct |
| c.7415A>C | p.(K2472T) | tcaagcagcagctgtaactttcacaacgtgtgaagaagaac | gttcttcttcacacgttgtgaaagttacagctgctgcttga |
| c.7544C>T | p.(T2515I) | ggcagtctgtatcttgcaaaaatatccactctgcc | ggcagagtggatatttttgcaagatacagactgcc |
| c.7879A>T | p.(I2627F) | ggtttataatcactatagatggttcatatggaaactggcagctat | atagctgccagtttccatatgaaccatctatagtgattataaacc |
| c.7940T>C | p.(L2647P) | atttgctaatagatgcccaagcccagaaagggtgc | gcaccctttctgggcttgggcatctattagcaaat |
| c.7958T>C | p.(L2653P) | ctaagcccagaaagggtgcctcttcaactaaaatacagat | atctgtattttagttgaagaggcaccctttctgggcttag |
| c.8063T>C | p.(L2688P) | cagctgcaaaaacacttgttccctgtgtttctgacataatttc | gaaattatgtcagaaacacagggaacaagtgtttttgcagctg |
| c.8165C>G | p.(T2722R) | cccaaaaagtggccattattgaacttagagatgggtggtatg | cataccacccatctctaagttcaataatggccactttttggg |
| c.8167G>C | p.(D2723H) | agtggccattattgaacttacacatgggtggtatgctg | cagcataccacccatgtgtaagttcaataatggccact |
| c.8187G>T | p.(K2729N) | agatgggtggtatgctgttaatgcccagttagatc | gatctaactgggcattaacagcataccacccatct |
| c.8243G>A | p.(G2748D) | gaatggcagactgacagttgatcagaagattattcttcatg | catgaagaataatcttctgatcaactgtcagtctgccattc |
| c.8525G>A | p.(R2842H) | atcatctggattatacatatttcacaatgaaagagaggaagaaaagg | ccttttcttcctctctttcattgtgaaatatgtataatccagatgat |
| c.8567A>C | p.(E2856A) | agcagcaaaatatgtggcggcccaacaaaagagac | gtctcttttgttgggccgccacatattttgctgct |
| c.9371A>T | p.(N3124I) | catatgttaattgctgcaagcatcctccagtggcg | cgccactggaggatgcttgcagcaattaacatatg |

**Table S5: *BRCA2* site-directed mutagenesis primers**. Complete list of all *BRCA2* mutagenesis primers used in this study. Human Genome Variant Nomenclature (HGVS) nucleotide and protein numbering shown for *BRCA2* reference sequence (GenBank: NM_000059.3).

| Nucleotide change | Amino acid change | Forward Mutagenesis Primer | Reverse Mutagenesis Primer |
|---|---|---|---|
| c.49_50delinsCC | p.(L17P) | cagctgtgaggagaaggaaaagccaaaggagaaattagcattcttg | caagaatgctaatttctcctttggcttttccttctcctcacagctg |
| c.53A>G | p.(K18R) | agctgtgaggagaaggaaaagttaagggagaaattagcattc | gaatgctaatttctcccttaacttttccttctcctcacagct |
| c.61_62delinsCC | p.(L21P) | gctgtgaggagaaggaaaagttaaaggagaaaccagcattcttgaaaaggg | cccttttcaagaatgctggtttctcctttaacttttccttctcctcacagc |
| c.70_71delinsCC | p.(L24P) | aaagttaaaggagaaattagcattcccgaaaagggaatacagcaagacacta | tagtgtcttgctgtattcccttttcgggaatgctaatttctcctttaacttt |
| c.71T>C | p.(L24S) | gttaaaggagaaattagcattctcgaaaagggaatacagcaagacac | gtgtcttgctgtattcccttttcgagaatgctaatttctcctttaac |
| c.83A>G | p.(Y28C) | agcattcttgaaaagggaatgcagcaagacactagccc | gggctagtgtcttgctgcattcccttttcaagaatgct |
| c.82_83delinsAC | p.(Y28P) | ggagaaattagcattcttgaaaagggaaaccagcaagacactagc | gctagtgtcttgctggtttcccttttcaagaatgctaatttctcc |
| c.90G>T | p.(K30N) | cttgaaaagggaatacagcaatacactagcccgcct | aggcgggctagtgtattgctgtattcccttttcaag |
| c.104T>C | p.(L35P) | cactagcccgccctcagcgtgccca | tgggcacgctgagggcgggctagtg |
| c.110G>A | p.(R37H) | gcccgccttcagcatgcccaaagagct | agctctttgggcatgctgaaggcgggc |
| c.110G>C | p.(R37P) | cccgccttcagcctgcccaaagagc | gctctttgggcaggctgaaggcggg |
| c.109C>A | p.(R37S) | agcccgccttcagagtgcccaaagagc | gctctttgggcactctgaaggcgggct |
| c.229delT | p.(C77Vfs*100) | gctaaaacactcagaacctaaaaataaaatagtgtttatgacaagttacacat | atgtgtaacttgtcataaacactattttatttttaggttctgagtgttttagc |
| c.557A>T | p.(N186I) | aggaacaggaagaaatcagtagcaaaattcctgctagatcac | gtgatctagcaggaattttgctactgatttcttcctgttcct |
| c.721A>G | p.(N241D) | cattcctaagaagacctgatttcaccagggcgact | agtcgccctggtgaaatcaggtcttcttaggaatg |
| c.925A>G | p.(I309V) | aacctccttgtaaataaagctgtaagtaaaagtggccaactgc | gcagttggccacttttacttacagctttatttacaaggaggtt |
| c.991G>C | p.(E331Q) | ttagaggcaaatatttcatgttctctaaatcaactcacctacaataac | gttattgtaggtgagttgatttagagaacatgaaatatttgcctctaa |
| c.1212T>A | p.(F404L) | gcctgaaggccttctgttacctgcagaatattatg | cataatattctgcaggtaacagaaggccttcaggc |
| c.1468C>G | p.(P490A) | tcattaactaaagtcagctctgccgctgggccc | gggcccagcggcagagctgactttagttaatga |
| c.1490delA | p.(N497Mfs*64) | gctgggcccactgaagatatgacttgtctagga | tcctagacaagtcatatcttcagtgggcccagc |
| c.1592delT | p.(L531Cfs*30) | gcatcagatcattgtgaaccactttgccaacttctagc | gctagaagttggcaaagtggttcacaatgatctgatgc |
| c.1676A>G | p.(Q559R) | atatcagcacgaaaaattatttattcgagtgaaagggaagaaaagtcg | cgacttttcttccctttcactcgaataaataatttttcgtgctgatat |
| c.2014G>C | p.(E672Q) | gatacagaaatggaggacttacaagaggaccttattgttcta | tagaacaataaggtcctcttgtaagtcctccatttctgtatc |
| c.2027T>C | p.(I676T) | aggacttagaagaggaccttactgttctaccaggaaaatc | gattttcctggtagaacagtaaggtcctcttctaagtcct |
| c.2134G>C | p.(A712P) | gtcattatcatcaggcggaaccgtatttaaaggagtataaagta | tactttatactcctttaaatacggttccgcctgatgataatgac |
| c.2135C>T | p.(A712V) | actcctttaaatacggttgtgcctgatgataatgacagg | cctgtcattatcatcaggcacaaccgtatttaaaggagt |
| c.2461A>T | p.(N821Y) | ccacccattgagtcattcacttttaaagaatatcagctctgtagaaa | tttctacagagctgatattctttaaaagtgaatgactcaatgggtgg |
| c.2590C>T | p.(P864S) | acctacaattggtttcagagttaaagaattcttcaggttcctgttc | gaacaggaacctgaagaattctttaactctgaaaccaattgtaggt |
| c.2794G>A | p.(V932M) | gcctgatgtgtataatctcatgtgtgtagctttgggaaa | tttcccaaagctacacacatgagattatacacatcaggc |
| c.2816T>G | p.(L939W) | ctcgtgtgtgtagctttgggaaattgggaaatcagagag | ctctctgatttcccaatttcccaaagctacacacacgag |
| c.2993G>A | p.(G998E) | gacgtttgcagaagatgaaggaggcaaagaaaacc | ggttttctttgcctccttcatcttctgcaaacgtc |
| c.3073_3074delinsCG | p.(A1025R) | aggtccaagggatgcaagaacgtctgcttggtactac | gtagtaccaagcagacgttcttgcatcccttggacct |
| c.3073G>A | p.(A1025T) | gaggtccaagggatgcaagaaactctgcttggta | taccaagcagagtttcttgcatcccttggacctc |
| c.3089C>T | p.(T1030I) | gcaagaagctctgcttggtactattattatgaacaacattgttattt | aaataacaatgttgttcataataatagtaccaagcagagcttcttgc |
| c.3110T>C | p.(I1037T) | gctgaggtccaagggacgcaagaagctctg | cagagcttcttgcgtcccttggacctcagc |
| c.3209T>C | p.(L1070P) | ctattctgaaatggggcttccctttattgtcctgagtcatc | gatgactcaggacaataaagggaagccccatttcagaatag |
| c.3290C>G | p.(P1097R) | gctcattgtgattaaccgtaagacgactctcagcg | cgctgagagtcgtcttacggttaatcacaatgagc |

**Table S6: *PALB2* site-directed mutagenesis primers**. Complete list of all *PALB2* mutagenesis primers used in this study. Human Genome Variant Nomenclature (HGVS) nucleotide and protein numbering shown for *PALB2* reference (GenBank: NM_024675.3).

| PALB2 nucleotide change | PALB2 amino acid change | ClinVar ID | Prior class | TLR results HR/ NHEJ mean | %HR mean | Published HR results Boonen et al.[2] %HR efficiency | Wiltshire et al.[3] HR fold change | Rodrigue et al.[4] % of WT HR |
|---|---|---|---|---|---|---|---|---|
| c.49_50delinsCC | p.(L17P) | N/A | VUS | 0.369 | 0.364 | | | |
| c.53A>G | p.(K18R) | 126758 | VUS | 1.212 | 1.049 | 100.19 | | >75 |
| c.61_62delinsCC | p.(L21P) | N/A | VUS | 0.245 | 0.251 | | | |
| c.70_71delinsCC | p.(L24P) | N/A | VUS | 0.330 | 0.300 | | | |
| c.71T>C | p.(L24S) | 230588 | VUS | 0.469 | 0.455 | 20.67 | 1.7 | |
| c.83A>G | p.(Y28C) | 126774 | VUS | 0.767 | 0.738 | 32.92 | 4.8 | 25-40 |
| c.82_83delinsAC | p.(Y28P) | N/A | VUS | 0.599 | 0.569 | | | |
| c.90G>T | p.(K30N) | 126779 | VUS | 1.221 | 1.377 | | 4.6 | |
| c.104T>C | p.(L35P) | 657328 | VUS | 0.328 | 0.383 | 10.4 | 0.8 | 5 |
| c.110G>A | p.(R37H) | 126590 | VUS | 1.064 | 0.981 | 44.9 | 4.1 | 25-40 |
| c.110G>C | p.(R37P) | N/A | VUS | 0.785 | 0.758 | | | |
| c.109C>A | p.(R37S) | 185108 | VUS | 1.102 | 0.944 | | 4.5 | |
| c.229delT | p.(C77Vfs*100) | 126644 | P/LP | 0.721 | 0.726 | | | |
| c.557A>T | p.(N186I) | 141993 | VUS | 1.254 | 1.241 | | | |
| c.721A>G | p.(N241D) | 126765 | B/LB | 1.419 | 1.296 | | | |
| c.925A>G | p.(I309V) | 126780 | B/LB | 1.517 | 1.361 | | 5.8 | |
| c.991G>C | p.(E331Q) | 241574 | VUS | 1.562 | 1.432 | | | |
| c.1212T>A | p.(F404L) | 418394 | VUS | 1.251 | 1.141 | | | |
| c.1468C>G | p.(P490A) | 241531 | VUS | 1.317 | 1.271 | | | |
| c.1490delA | p.(N497Mfs*64) | 460897 | P/LP | 0.365 | 0.317 | | | |
| c.1592delT | p.(L531Cfs*30) | 126609 | P/LP | 0.285 | 0.309 | 7.75 | | |
| c.1676A>G | p.(Q559R) | 126613 | B/LB | 1.300 | 1.143 | 95.02 | | |
| c.2014G>C | p.(E672Q) | 126630 | B/LB | 1.083 | 0.950 | 79.52 | | |
| c.2027T>C | p.(I676T) | 142310 | B/LB | 1.214 | 1.137 | | | |
| c.2134G>C | p.(A712P) | 186485 | VUS | 1.542 | 1.559 | | | |
| c.2135C>T | p.(A712V) | 126637 | B/LB | 1.422 | 1.234 | | | |
| c.2461A>T | p.(N821Y) | 480228 | VUS | 1.094 | 1.008 | | | |
| c.2590C>T | p.(P864S) | 126669 | B/LB | 1.073 | 1.066 | 85.8 | | >75 |
| c.2794G>A | p.(V932M) | 126682 | B/LB | 1.188 | 1.143 | | | >75 |
| c.2816T>G | p.(L939W) | 126683 | VUS | 1.090 | 0.975 | 60.28 | 4.8 | |
| c.2993G>A | p.(G998E) | 126699 | B/LB | 1.348 | 1.206 | 95.16 | | >75 |
| c.3073_3074delinsCG | p.(A1025R) | N/A | VUS | 0.479 | 0.430 | 17.62 | | |
| c.3073G>A | p.(A1025T) | 217917 | VUS | 1.096 | 0.995 | | 3.9 | |
| c.3089C>T | p.(T1030I) | 232977 | VUS | 0.859 | 0.824 | 14.68 | 3 | 23.6 |
| c.3110T>C | p.(I1037T) | 484219 | VUS | 1.230 | 1.212 | 38.86 | | |
| c.3209T>C | p.(L1070P) | 216752 | VUS | 0.641 | 0.596 | 23.09 | 1.7 | |
| c.3290C>G | p.(P1097R) | 140834 | VUS | 0.308 | 0.298 | | | |
| Empty vector | Empty vector | N/A | Control | 0.568 | 0.503 | | | 4.38 |
| Wild-type | Wild-type | N/A | Control | 1.284 | 1.221 | | | 100 |

**Table S7: TLR performance compared to recent publications.** Comparison of TLR data to published *PALB2* results, as in Figure 5. Clinical Variant Database (ClinVar)[5] accession numbers provided as available. Normalized TLR results shown as mean of at least 3 replicates. For Boonen et al.[2], 5-10% of WT HR efficiency was considered abnormal and 10-40% was considered intermediate function. Wiltshire et al.[3] considered scores of 1.7 or less (34% of WT) to be abnormal and set 2.4 (48% of WT) as the upper threshold for intermediate function. Below 25% and 40% of WT HR activity was considered abnormal and intermediate function, respectively, by Rodrigue et al.[4] Key: B/LB, benign/likely benign; HR, homologous recombination; NHEJ, non-homologous end-joining; P/LP, pathogenic/likely pathogenic; TLR, traffic light reporter; VUS, variant of uncertain significance; WT, wild-type. HGVS numbering shown for *PALB2* reference (GenBank: NM_024675.3).
